# STAGdb: a 30K SNP genotyping array and Science Gateway for Acropora corals and their dinoflagellate symbionts

**DOI:** 10.1101/2020.01.21.914424

**Authors:** S.A. Kitchen, G. Von Kuster, K.L. Vasquez Kuntz, H.G. Reich, W. Miller, S. Griffin, Nicole D. Fogarty, I.B. Baums

**Author notes:** Corresponding Author: Iliana B. Baums, The Pennsylvania State University 208 Mueller Laboratory University Park, PA, 16802.

## Abstract

Standardized identification of genotypes is necessary in animals that reproduce asexually and form large clonal populations such as coral. We developed a high-resolution hybridization-based genotype array coupled with an analysis workflow and database for the most speciose genus of coral, *Acropora*, and their symbionts. We designed the array to co-analyze host and symbionts based on bi-allelic single nucleotide polymorphisms (SNP) markers identified from genomic data of the two Caribbean *Acropora* species as well as their dominant dinoflagellate symbiont, *Symbiodinium ‘fitti’.* SNPs were selected to resolve multi-locus genotypes of host (called genets) and symbionts (called strains), distinguish host populations and determine ancestry of the coral hybrids in Caribbean acroporids. Pacific acroporids can also be genotyped using a subset of the SNP loci and additional markers enable the detection of symbionts belonging to the genera *Breviolum, Cladocopium*, and *Durusdinium*. Analytic tools to produce multi-locus genotypes of hosts based on these SNP markers were combined in a workflow called the <u>S</u>tandard <u>T</u>ools for <u>A</u>croporid <u>G</u>enotyping (STAG). In the workflow the user’s data is compared to the database of previously genotyped samples and generates a report of genet identification. The STAG workflow and database are contained within a customized Galaxy environment (https://coralsnp.science.psu.edu/galaxy/), which allows for consistent identification of host genet and symbiont strains and serves as a template for the development of arrays for additional coral genera. STAG data can be used to track temporal and spatial changes of sampled genets necessary for restoration planning as well as be applied to downstream genomic analyses.

## Introduction

Genotype identification and tracking are required for well-replicated basic research experiments and in applied research such as designing restoration projects. High-resolution genetic tools are necessary for large clonal populations where genets can only be delineated via genotyping. The advent of reduced representation sequencing methods such as Genotype-By-Sequencing (GBS) or Restriction-site Associated DNA Sequencing (RADseq) have made it possible to assay a large number of single-nucleotide polymorphism (SNP) loci in any organism at a reasonable cost (Altshuler et al. 2000). These methods are widely used in population genomics but have the disadvantage that the SNP loci are anonymous. Thus, there is no guarantee that the same set of SNP loci will be recovered from each sample within an experiment or between experiments, making it more difficult to design standardized workflows. To circumvent this issue, standardized SNP probes can be designed for reproducible genotyping and analysis from hundreds of samples using modified RAD-based approaches like Rapture (Ali et al. 2016), RADcap (Hoffberg et al. 2016), and quaddRAD (Franchini et al. 2017) or using hybridization-based SNP genotyping arrays. Hybridization-based SNP arrays tend to have lower error rates then RADseq methods (Darrier et al. 2019; Palti et al. 2015) and thus increased accuracy of genet identification and tracking. However, both approaches forgo discovery of new SNP loci in favor of assaying a standard set of probes across all samples resulting in some ascertainment bias (Moragues et al. 2010; Malomane et al. 2018; Lachance and Tishkoff 2013).

When it comes to the analysis of SNP genotyping data, familiarity with computer programming and access to high performance computing is typically required but not always available. Because genotyping arrays contain a known set of SNP loci, standardized workflows can be designed easily. Galaxy is an open source, web-based platform for data-intensive biomedical research (Afgan et al. 2018) and provides the underlying framework for Science Gateways. Science Gateways are extensions of cyberinfrastructure, like Galaxy, that focus on a specific scientific communities’ needs by providing digital interfaces of computational resources which lowers the barriers (know-how and cost) often associated with these resources. The use of a standardized workflow within a Scientific Gateway enables scientists and restoration practitioners to accurately match samples to existing genets and strains, discover novel genets/strains and track their fate across years, all from a web browser.

Corals, like other clonal plant and animal species, reproduce frequently via asexual fragmentation (Whitaker 2006; Miller and Ayre 2004; Stoddart 1983; Ayre and Hughes 2000; Adjeroud and Tsuchiya 1999). Over time coral genets can extend over tens of meters consisting of tens to hundreds of colonies (Foster et al. 2007; Neigel and Avise 1983; Baums et al. 2006). This leads to considerable variability in genotypic evenness and richness on small spatial scales, ranging from minimal clonal replication to reefs dominated by a single genet (Pinzón et al. 2012; Baums et al. 2006; Fig 1, Ayre and Hughes 2000; Miller and Ayre 2004). The importance of coral genets in explaining variation in growth rates and stress response is becoming increasingly clear (Parkinson and Baums 2014; Polato et al. 2013; Baums et al. 2013; Randall and Szmant 2009; Meyer et al. 2009). Further, hermaphroditic corals species like the Caribbean acroporids are mainly self-incompatible, thereby requiring the presence of gametes from different genets for successful sexual reproduction (Baums et al. 2005a; Fogarty et al. 2012). For these reasons, identification of genets and preservation of genotypic diversity are conservation priorities (Baums et al. 2019).

**Figure 1.**
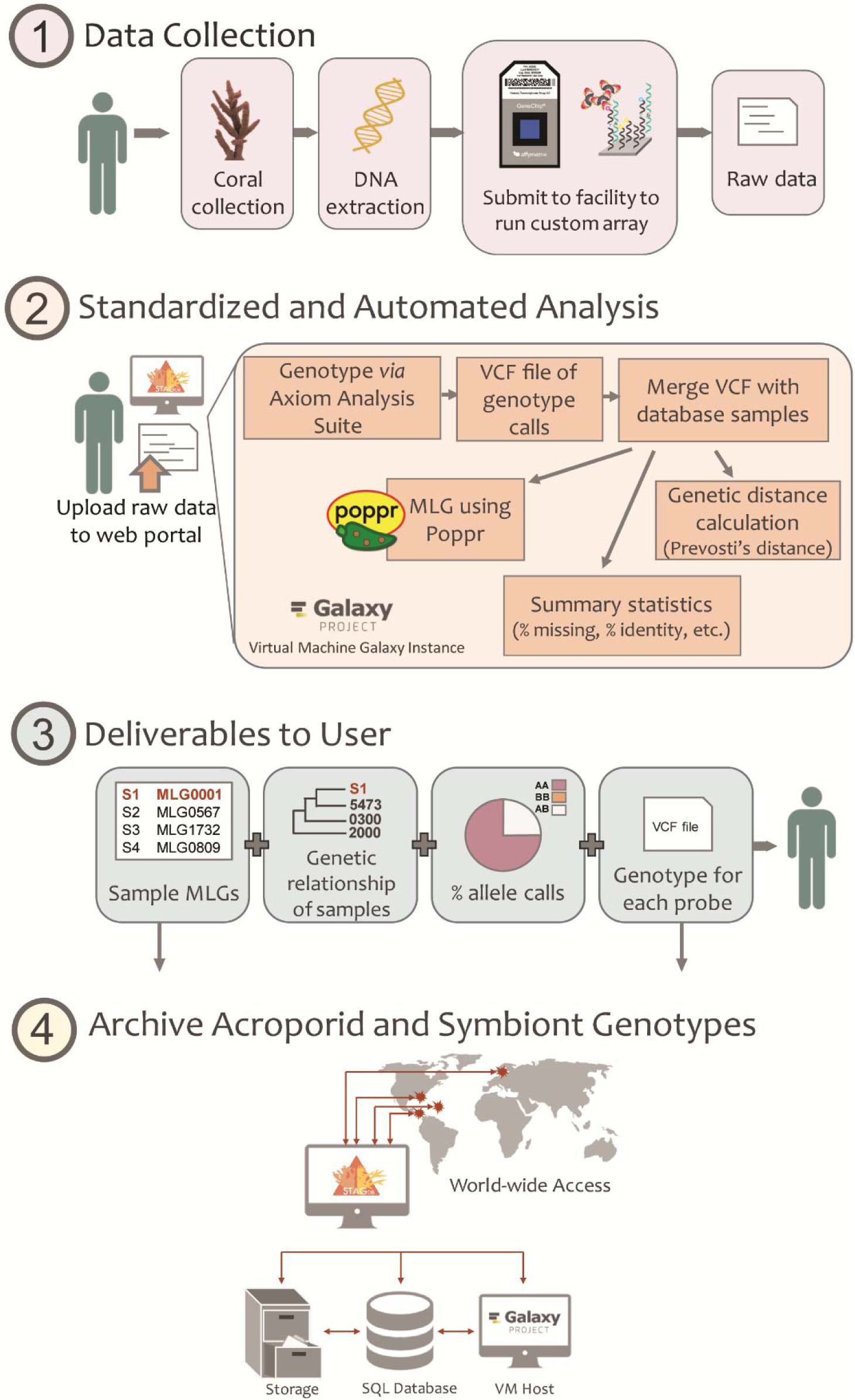
General overview of Standard Tools for Acroporid Genotyping. Step 1) user collects the coral, extracts the DNA and submits the DNA to their closest processing facility. Step 2) user uploads metadata and raw data to the Galaxy CoralSNP environment for analysis. Step 3) user downloads their multi-locus genotypes (MLG) among other deliverables. Step 4) the new sample MLGs and genotype information is deposited in the postgreSQL database that can be accessed from anywhere.

Tropical corals frequently house single-celled photosynthetic algae in the family Symbiodiniaceae that provide the majority of the hosts organic carbon (Muscatine and Cernichiari 1969; Davies 1991). Coral species differ in their symbiont specificity, and colonies may house several algal genera within their cells at a given time. Thus, the complex mixtures of coral and algal DNA present challenges and opportunities for the development of high-resolution co-genotyping methods. Microsatellite markers specific for certain species of algae have further revealed subspecies level strain diversity and elucidated the temporal and spatial dynamics of symbiont strain/host genet associations (Santos and Coffroth 2003; Pettay and LaJeunesse 2007; Pettay and LaJeunesse 2009; Pinzón et al. 2011; Baums et al. 2014; Wham et al. 2011; Grupstra et al. 2017; Chan et al. 2019; Andras et al. 2009), but no SNP-based markers are available yet.

Given that the algal species associated with a coral colony can influence the colony’s physiology, it is also of interest to researchers and practitioners to identify the dominant and any background symbionts in coral samples.

Corals often occur in remote locations without access to molecular laboratory and computation facilities, or require special export permits to transport tissue samples to well-equipped facilities. Thus, we aimed to develop a genotyping array designed for instruments available at most major hospitals around the world. Genotyping arrays can be processed by a sequencing facility with user supplied tissue (as well as extracted DNA; Figure 1) eliminating the need for a molecular laboratory and therefore, can be widely adopted by users without access to such facilities.

Here, we report the development of a SNP array and standardized analysis workflow for the most speciose genus of coral, *Acropora*. The roughly 120 *Acropora* spp. dominate shallow reefs in the Pacific and Atlantic oceans (Veron 2000; Wallace 1999). In the Caribbean, the primary shallow reef builders are *Acropora palmata* and *A. cervicornis,* which form a hybrid (commonly known as *A. prolifera*) (van Oppen et al. 2000; Vollmer and Palumbi 2002a; Lamarck 1816). Because of drastic population declines, they are listed as threatened under the U.S. Endangered species act, making them the focal species in reef restoration efforts across the Caribbean. Promoting genotypic diversity within nurseries and outplanting sites is a management priority for these species. We present a ∼30k SNP genotyping array that identifies host and symbiont genotypes, coral hybrid status and background symbiont genera. The array can be analyzed cost-effectively in a standardized manner using the Standard Tools for Acroporid Genotyping (STAG) within a Galaxy environment (Figure 1). We further establish a publicly available database of *Acropora* genets. This approach can serve as a template for other asexually producing species of conservation concern.

## Methods and Materials

### Coral collection and DNA extraction

Initially samples were selected from an archival tissue collection to address technical concerns regarding: dual genotyping with enrichment of host or symbiont DNA, tissue type (sperm, larvae and adult), clone identification (multiple ramets per genet), reproducibility between labs (Baums lab and Dr. Nicole Fogarty’s lab), and difference in tissue preservatives (flash frozen, non-denatured 95-100% ethanol, CHAOS buffer and DMSO). We also included several *Acropora* samples from the Pacific including *A. muricata*, *A. millepora* and *A. digitifera* (generously donated by Drs. Todd LaJeunesse, Zachary Fuller, Mikhail Matz and Stephen Palumbi). The remaining samples include archival samples collected from various locations across the geographic distribution of *A. palmata* and *A. cervicornis* and their hybrid, *A. prolifera*. Sample information can be found in Supplemental Table 1.

DNA extraction methods varied depending on the tissue type, sample preservative and laboratory as follows (also see Table S1). High molecular weight DNA from concentrated coral sperm was extracted using the illustra Nucleon Phytopure kit (GE Healthcare Life Science, Pittsburgh, PA) following the manufacturer’s instructions and eluted in nuclease-free water. For coral larvae, an individual larva was incubated in 12 µl of lysis solution (10.8 µl Buffer TL (Omega BioTek, Norcross, GA), 1 µl of OB Protease Solution (Omega BioTek) and 0.2 µl of RNAse A (100 mg/ml)) for 20 min at 55 °C. An additional 38 µl of Buffer TL was added to each sample followed by 50 µl of phenol/chloroform/isoamyl alcohol solution (25:24:1) and gently rocked for 2 min. The samples were centrifuged for 10 min at 10,000 rpm. To the aqueous phase, 50 µl of chloroform:isoamyl alcohol (24:1) was added and mixed with gentle rocking for 2 min. The aqueous phase was recovered after 5 min centrifugation at 10,000 rpm. The DNA was precipitated with 1.5x volume of room-temperature isopropanol, 1/10 volume of 3M sodium acetate (pH=5.2) and 1 µl of glycogen (5 mg/ml) for 10 min at room temperature followed by centrifugation at 15,000 rpm for 20 min and two rounds of washes with 70% ethanol. The pellets were resuspended in 20 µl of low TE buffer (10 mM Tris-HCl and 0.1 mM EDTA). In the Baums lab, DNA from adult coral tissue was extracted using the Qiagen DNeasy kit (Qiagen, Valencia, CA) following the manufacturer’s protocol and eluted in 100 µl of nuclease-free water or low TE buffer. A subset of adult tissue was extracted separately by Dr. Fogarty’s lab. These samples were either preserved directly in CHAOS DNA extraction buffer (Fukami et al. 2004) or ethanol and the DNA was isolated using a magnetic bead protocol (Levitan et al. 2011). These methods are referred to as a “mixed” extraction because both coral and symbiont DNA is recovered, but in unknown proportions. For enriched symbiont DNA from the coral tissue, we isolated the symbionts using a modification of Wayne’s method (Wilson et al. 2002) described by Bongaerts et al. (2017). Briefly, ∼3-4 coral calyces were placed in 600 µl of nuclease-free water and vortexed on maximum speed for 1 min to remove the tissue from the coral skeleton.

**Table 1.** Number of recommended probes for each taxon from five independent runs.

|  |  | Caribbean *Acropora* |  |  |  | Symbiont |  |  |  |
| --- | --- | --- | --- | --- | --- | --- | --- | --- | --- |
| Plate Number | Probe Classification | Fixed | Population | Variable | Total | Fixed | Population | Genera | Total |
| All probes |  | 25,889 | 17,803 | 9,887 | 53,579 | 3,663 | 304 | 54 | 4,021 |
| Plate 1 | Recommended | 9,919 | 6,455 | 3,342 | 19,716 | 2,176 | 109 | 18 | 2,303 |
|  | polyHigh | 9,663 | 5,855 | 2,515 | 18,033 | 2,093 | 68 | 0 | 2,161 |
|  | MonoHigh | 181 | 173 | 524 | 878 | 83 | 41 | 14 | 138 |
|  | noMinor | 75 | 427 | 303 | 805 | 0 | 0 | 0 | 0 |
| Plate 2 | Recommended | 10,199 | 6,701 | 3,445 | 20,345 | 2,210 | 100 | 15 | 2,325 |
|  | polyHigh | 9,987 | 6,381 | 2,759 | 19,127 | 1,792 | 68 | 1 | 1,861 |
|  | MonoHigh | 165 | 164 | 572 | 901 | 418 | 32 | 12 | 462 |
|  | noMinor | 47 | 156 | 114 | 317 | 0 | 0 | 2 | 2 |
| Plate 3 | Recommended | 12,299 | 7,823 | 4,137 | 24,259 | 2,189 | 109 | 17 | 2,315 |
|  | polyHigh | 11,916 | 6,918 | 3,249 | 22,083 | 261 | 54 | 0 | 315 |
|  | MonoHigh | 195 | 189 | 681 | 1,065 | 1,928 | 55 | 14 | 1,997 |
|  | noMinor | 188 | 716 | 207 | 1,111 | 0 | 0 | 3 | 3 |
| Plate 4 | Recommended | 10,171 | 7,178 | 3,814 | 21,163 | 2,247 | 107 | 16 | 2,370 |
|  | polyHigh | 8,989 | 5,783 | 2,827 | 17,599 | 223 | 48 | 0 | 271 |
|  | MonoHigh | 435 | 593 | 774 | 1,802 | 2,024 | 59 | 16 | 2,099 |
|  | noMinor | 746 | 802 | 213 | 1,761 | 0 | 0 | 0 | 0 |
| Plate 5 | Recommended | 11,093 | 7,620 | 4,038 | 22,751 | 2,215 | 113 | 17 | 2,345 |
|  | polyHigh | 8,748 | 5,662 | 2,498 | 16,908 | 621 | 37 | 1 | 659 |
|  | MonoHigh | 459 | 580 | 821 | 1,860 | 1,594 | 76 | 16 | 1,686 |
|  | noMinor | 1,886 | 1,378 | 719 | 3,983 | 0 | 0 | 0 | 0 |

The skeleton pieces were removed and the supernatant with the coral tissue was centrifuged for 3 min at 2,500 rpm. The supernatant was removed leaving a pellet of symbiont cells. Glass beads were added to the symbiont pellet and vortexed on high for 1 min to disrupt the cell membrane. High-molecular weight DNA quality was assessed using gel electrophoresis, and yield quantified using either NanoDrop 2000 (Thermo Scientific) or PicoGreen assay (Invitrogen) for sperm and adult extractions and a Qubit fluorometer dsDNA Broad Range kit (Invitrogen) for the larval extractions.

### Coral SNP selection

We previously identified 4.9 million SNPs between the two Caribbean acroporids and the Pacific acroporid *A. digitifera,* and of those 1.6 million high-quality SNPs varied between the Caribbean acroporids (Kitchen et al. 2019). To create a conservative set of SNP, we additionally called variants with freebayes v1.1.0-50-g61527c5 (Garrison and Marth 2012) using the same alignment file from the previous study and identified shared SNPs between the two variant callers with vcf-compare v0.1.14-12-gcdb80b8 (Danecek et al. 2011). From these shared SNPs, they were further refined into three informative categories: fixed, population and variable (Figure 2B). The “fixed” SNPs are those variants where all 21 individuals of a given species share a nucleotide and the other 21 individuals of the other species share a different nucleotide. The fixed SNPs were filtered to a sample read depth of ≥ 3 and a minimum distance of 500 bp. We also retained those that we previously defined as PCR-ready (*n*= 894, no observed SNPs, indels, low-complexity DNA or unassembled regions within 50 bp on either side of the SNP, see (Kitchen et al. 2019). Population SNPs were identified based on pairwise comparisons of the four different collection sites (Table S2). These SNPs were filtered such that all samples from one site shared an allele with a frequency of 0.8 or greater and differed from the samples of the other site with the alternative allele at a frequency of 0.8 or greater. Finally, variable SNPs were identified by filtering the SNPs to a sample read depth of ≥ 4, allowing no ambiguous bases or repetitive sequences in 71 bp of flanking sequence, a minimum distance of at least 1,000 bp between surrounding SNPs, and an allele frequency between 0.5 and 0.7 for all 21 *A. palmata* samples while the variants was also observed in the *A. cervicornis* samples. SNP frequencies were calculated using –freq parameter with VCFtools (Danecek et al. 2011).

**Figure 2.**
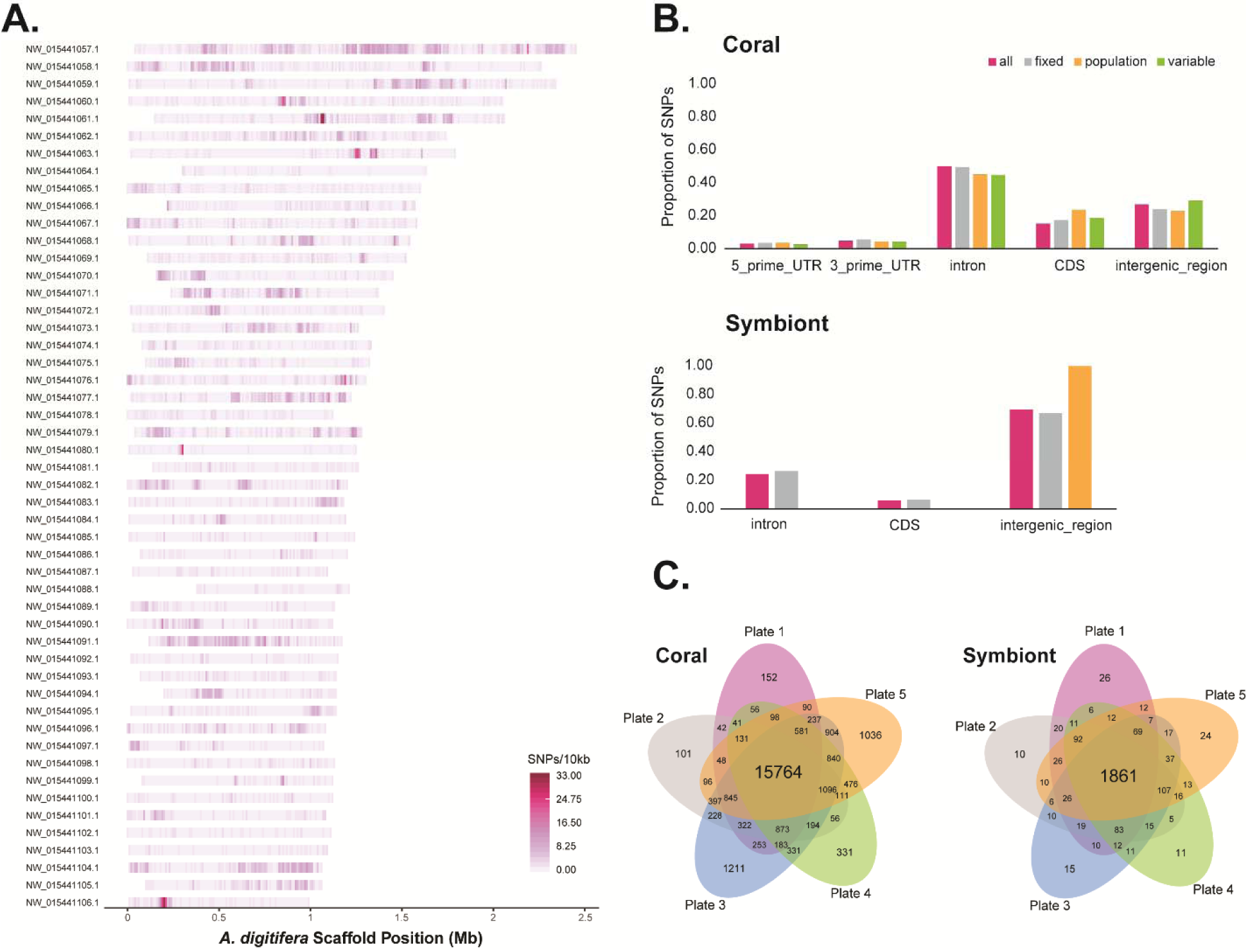
Density, distribution and recovery of SNP probes. The probe density over 10,000 bp windows is mapped onto the 50 longest *A. digitifera* reference scaffolds (A). The highest density exceeds 33 probes in a given interval, where most intervals are between 0 to 8 probes. The proportion of designed probes are compared for coding and non-coding regions in the genomes of the coral and symbionts (B). All probes are pink, fixed probes are grey, population probes are orange, and variable probes are green. The recommended probes shared between each plate are shown in the Venn diagrams (C).

**Table 2.** Success rate after quality-filtering of Caribbean acroporid and symbiont samples. To the left of the slash are those samples that passed the default quality filtering of the Best Practices Workflow (BPW) and to the right is the total number of processed samples.

|  | Plate 1 |  | Plate 2 |  | Plate 3 |  | Plate 4 |  | Plate 5 |  |
| --- | --- | --- | --- | --- | --- | --- | --- | --- | --- | --- |
|  | Coral | Symbiont | Coral | Symbiont | Coral | Symbiont | Coral | Symbiont | Coral | Symbiont |
| **All samples** | 90/95 | 81/90* | 92/96 | 78/95* | 93/95 | 29/84 | 72/96 | 23/72* | 90/96 | 70/96* |
| *Mixed extractions* | 81/84 | 76/89* | 91/95 | 78/95* | 83/83 | 29/83 | 72/72 | 23/72* | 90/96 | 70/96* |
| *Symbiont-enriched extractions* | 3/5 | 5/5 | -- | -- | -- | -- | -- | -- | -- | -- |
| *Sperm* | 1/1 | -- | 1/1 | -- | -- | -- | -- | -- | -- | -- |
| *Larvae* | 5/5 | -- | -- | -- | 10/12 | -- | -- | -- | -- | -- |
| *Symbiont culture* | -- | 0/1* | -- | -- | -- | 0/1* | -- | -- | -- | -- |
| *Indo-Pacific acroporid* | 0/1* | -- | -- | -- | -- | -- | 0/24* | -- | -- | -- |

\* Failed on dish quality (signal observed from non-polymorphic loci) – indication of different coral/symbiont species or background symbionts.

For each SNP, 35 bp of identical flanking sequence between the species was pulled from the *A. digitifera* genome assembly (NCBI: GCF_000222465.1; Shinzato et al. (2011)) using bedtools getfasta (Quinlan and Hall 2010). These 71 nucleotide (71mer) candidate sequences were filtered through a series of similarity searches to reduce non-specific sequence capture. First, the sequences were compared to the *A. digitifera* genome assembly using BLAST v2.6.0 (task= blastn, e-value= 1e-13) to determine whether redundant genomic targets were present. Sequences were discarded that had a ≥ 30 bp match with more than one genomic location. To check for repetitive probes, a same-strand self-analysis was performed using blastn (filter query sequence = false, word size =11, -dust no, e-value = 1e-13, strand = both).

In addition to the SNP probes, we identified non-polymorphic sequences from acroporids by extracting high-quality SNPs that were identical between the two Caribbean acroporids and different from *A. digitifera*. We required a sample read depth of ≥ 6 reads with a minimum distance of 1,000 bp between SNPs and no repetitive or ambiguous bases in the 35 bp flanking sequence. We searched for sequence similarity between these probes and the final acroporid probe sets using BLAST (task= blastn, e-value= 1e-13). We discarded those that had significant overlap to the array probes and randomly selected 3,000 to act as the background probes.

### Symbiont variant calling and SNP selection

SNP discovery in the symbionts was accomplished by comparing our genome samples to two reference genomes, either the assembly of cultured *S. tridacnidorum* (NCBI: GCA_003297005.1, (Shoguchi et al. 2018)) or partial assembly of the predominant symbiont of *A. palmata* and *A. cervicornis*, *S. ‘fitti’* (Reich et al., unpublished), both of which belong to the genus *Symbiodinium* (ITS2-clade A3). Because the genome samples were not specifically extracted for symbiont DNA, only 15-25% of the reads mapped to the symbiont genomes reducing our ability to identify comparable number of SNPs in the symbiont as the coral. Some of the *Symbiodinium* SNPs were identified by comparing only the deep-coverage metagenome sequences of *A. palmata* and *A. cervicornis* to the genome assembly of *S. tridacnidorum* (Shoguchi et al. 2018). These SNPs were identified as fixed between the two representative Florida acroporids sampled, but it was unclear if they were fixed between the symbiont strains of the two coral species across their range, just in Florida or just between these two samples. The other *Symbiodinium* SNPs were identified by mapping the 42 re-sequenced genome samples to a draft genome assembly of *S. ‘fitti’* and processed as described in (Kitchen et al. 2019). High-quality SNPs were those with quality phred score > 200 and no more than 20% missing data at a given site among all samples. The 71 bp flanking sequences were filtered through the series of blast homology searches in the same manner as the coral SNPs described above. Finally, to confirm that the probes designed for the host and symbiont did not overlap, the final set of both groups were compared to each other using blastn with an e-value threshold of 1e-13. No matches were found between the acroporid and symbiont probe sequences. For the *Symbiodinium* non-polymorphic SNVs, we extracted genomic regions from the *S. ‘fitti’* scaffolds with the highest gene coverage for the *A. palmata* and *A. cervicornis* shallow genome samples. After searching them against each other using blastn (task= blastn, e-value= 1e-13), a random subset of 3,000 probes was selected.

In addition to the genotyping probes, we identified 12 SNPs in loci used to distinguish genera of Symbiodinaceae to capture potential background symbionts. The most common genera associated with tropical corals are *Symbiodinium*, *Breviolum*, *Cladocopium* and *Durisdinium* and can be distinguished by genetic markers. These loci include ribosomal (*internal transcribed spacer 2* and *nr28S*), mitochondrial (*COI* and *cob*), chloroplast (*cp23S* and *psbA*) and nuclear (*elongation factor 2*) markers using sequences from previously published studies (Takishita et al. 2003; Pochon et al. 2012; Arif et al. 2014; LaJeunesse 2002; LaJeunesse 2001). Sequence accessions are provided in Supplemental Table S3. At least one representative sequence from each of the genera *Symbiodinium*, *Breviolum*, *Cladocopium* and *Durusdinium* for each locus was aligned with MUSCLE in Mega X (Kumar et al. 2018). SNPs were identified based on their ability to distinguish genera with enough conserved flanking sequence for probe design (Table S4).

**Table 3.** Summary of population genetic variation of Caribbean acroporids estimated with 19,694 genotyping probes.

| Species | Population | N | N<sub>G</sub> | H<sub>O</sub> | H<sub>S</sub> | F<sub>IS</sub> |
| --- | --- | --- | --- | --- | --- | --- |
| *A. cervicornis* | Belize | 27 | 18 | 0.117 | 0.122 | 0.033 |
|  | Cuba | 1 | 1 | NA | NA | NA |
|  | Curacao | 9 | 7 | 0.126 | 0.128 | -0.002 |
|  | Florida | 54 | 46 | 0.110 | 0.113 | 0.038 |
|  | Puerto Rico | 35 | 21 | 0.127 | 0.118 | -0.042 |
|  | USVI | 16 | 9 | 0.113 | 0.109 | -0.023 |
| *A. palmata* | Belize | 37 | 27 | 0.148 | 0.149 | 0.013 |
|  | Curacao | 73 | 57 | 0.132 | 0.124 | 0.013 |
|  | Florida | 58 | 26 | 0.151 | 0.154 | 0.021 |
|  | Puerto Rico | 132 | 75 | 0.141 | 0.140 | 0.000 |
|  | USVI | 8 | 8 | 0.156 | 0.156 | -0.004 |
| *A. prolifera* | Antigua | 8 | 8 | 0.656 | 0.410 | -0.543 |
|  | Bahamas | 2 | 2 | 0.692 | 0.415 | -0.705 |
|  | Belize | 21 | 21 | 0.674 | 0.406 | -0.580 |
|  | Cuba | 2 | 2 | 0.679 | 0.415 | -0.673 |
|  | Curacao | 4 | 4 | 0.689 | 0.382 | -0.770 |
|  | USVI | 2 | 2 | 0.700 | 0.412 | -0.725 |

N= number of samples, N<sub>G</sub>= number of genets, H<sub>O</sub>= average observed heterozygosity, H<sub>S</sub>= average expected proportion of heterozygote individuals in the subpopulations, and F<sub>IS</sub>= average inbreeding coefficient.

### SNP validation by genotyping

After filtering, 34,783 acroporid SNPs (15,644 fixed, 10,429 population and 6,050 variable) and 2,661 symbiont SNPs were submitted for review by Affymetrix (Thermo Fisher, Santa Clarita, CA). Final probe construction was completed by their bioinformatics team (Table 1). Every SNP locus submitted had at least one probe on the final array. For a set of high-priority SNPs, at least two probes were designed and then the remainder of the array was filled with the highest scoring SNPs until full. The final coral probe set was run through snpEff v4.3 (Cingolani et al. 2012) and the final algal probe set was compared to the respective GFF file for each *Symbiodinium* genome using bedtools intersect (Quinlan and Hall 2010) to determine genomic location. The SNP density in bin sizes of 10,000 was extracted for all coral probes using VCFtools v0.1.15 (Danecek et al. 2011).

We worked with Affymetrix to optimize their current genotyping tools and pipeline to provide dual genotyping of the coral and symbiont in a single run. Five 96-well plates were submitted to the Affymetrix team for processing the Axiom Mini 96 custom genotype array on the GeneTitan (Thermo Fisher, Santa Clarita, CA). The raw data was analyzed using the Axiom ‘Best Practices Workflow’ (BPW) in the Axiom Analysis Suite software (Thermo Fisher, Santa Clarita, CA) for each of the five runs separately for the coral and algal probe sets, with default quality filtering thresholds. Important thresholds that identify low sample quality include the dish quality, which is the signal of the non-polymorphic probes from one individual to the next, and call rate, which is the proportion of assigned genotypes for an individual out of all tested probes. The Bayesian clustering algorithm BRLMM-P (Affymetrix 2007) was used to compute three posterior cluster locations (AA, AB, and BB) based on pre-positioned genotype cluster locations called priors. Genotype calls were made by identifying the intensity distribution, or cluster, each sample most likely belongs to with a confidence score (1 – posterior probability of the sample assignment to genotype cluster). In the case of the symbionts, because they are haploid, the algal genotyping probes were treated as mitochondrial probes with only homozygous AA or BB allele calls being valid. Five of the symbiont genera probes allowed for three clusters when multiple alleles were predicted to separate different genera (Table S4).

Following the analysis of the five plates, the performance of each probe was classified into six categories based on their separation of genotype clusters with SNPpolisher (Affymetrix, CA) (Table 1). These categories include Poly High Resolution, Mono High Resolution, No Minor Hom, Call Rate Below Threshold, Other and Off-Target Variant. Probes that fell under ‘Poly High Resolution’ are those with resolution of three clusters (AA, AB and BB) with at least two sample having the minor allele. Probes that fell under ‘Mono High Resolution’ are those where all samples share the same allele possibly due to low minor allele frequency or sample selection on the plate. Finally, probes that fell under ‘No Minor Hom’ are those where no minor homozygous allele is observed, only AA and AB. These three categories make up the “best and recommended” probe set that was used in downstream analyses (Table S5-S7, and Figure 2C).

### Standard Tools for Acroporid Genotyping workflow

The general overview of the data conversion and genotype analysis steps are presented in Figure 3A and code for new Galaxy tools can be found at (https://github.com/gregvonkuster/galaxy_tools/tree/master/tools/corals). The resulting genotype files from the BPW were first converted into the variant call format (VCF) using the bcftools plugin affy2vcf (https://github.com/freeseek/gtc2vcf). The VCF files were sorted and merged using bcftools (Figure 3A). The coral genotyping probes recommended from the BPW with the first plate were subset from the VCF of previous genome samples using VCFtools. These SNPs were marked in the INFO field that was then used for filtering after the array data was merged with the genome samples (Figure 3A, “Select” step). The Affymetrix IDs and the order of the samples were extracted from the filtered VCF using the *Affy Ids for Genotyping* tool and combined with the sample attributes from the BPW and population information in the user-supplied metadata into a new text file with the *Genotype Population Info* tool (Figure 3A). Additional population information was appended to the file from the previously genotyped samples in the database.

**Figure 3.**
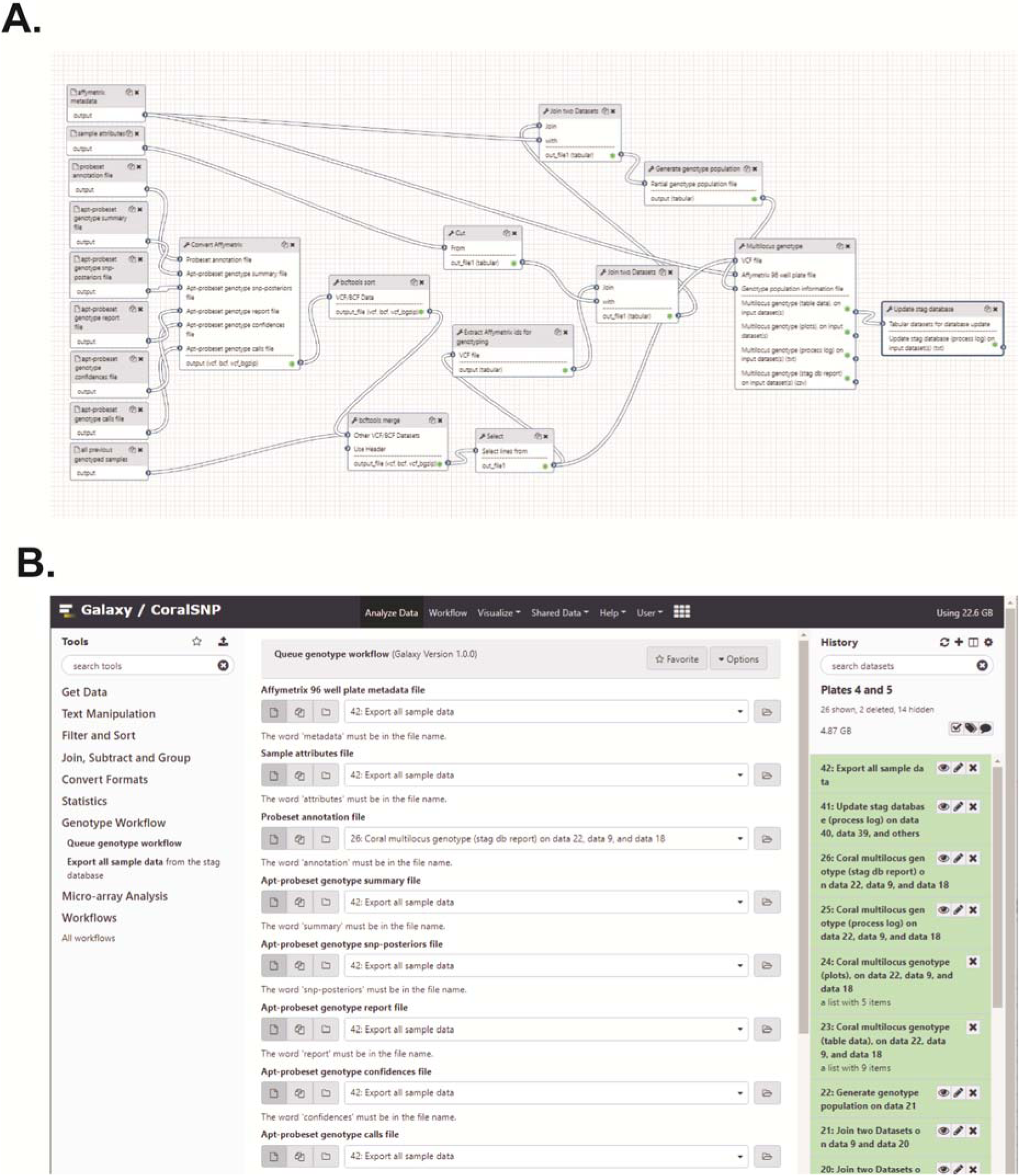
Galaxy CoralSNP workflow and interface. This analysis pipeline (A) is initiated by the *Queue Genotype Workflow* tool via the Galaxy REST API. The workflow consists of the following tools, all of which can be installed into a local Galaxy instance from the Main Galaxy Tool Shed (https://toolshed.g2.bx.psu.edu). Affy2vcf2 converts Affymetrix genotype calls and intensity files to the VCF format. Bcftools_sort sorts bcf/vcf files. Bcftools_merge merges bcf/vcf files. Affy_ids_for_genotyping extracts information from a VCF files that contains Affymetrix identifiers and produces a file that contains a subset of the identifiers combined with additional data to generate the genotype population information for use as input to the *Coral Multilocus Genotype* tool. Genotype_population_info generates the genotype population information file for use as input to the *Coral Multilocus Genotype* tool. Coral_multilocus_genotype renders the unique combination of alleles for two or more loci for each individual. Update_stag_database updates the stag database tables from a dataset collection where each item in the collection is a tabular file that will be parsed to insert rows into a table defined by the name of the file. The code for these tools is available in GitHub at https://github.com/gregvonkuster/galaxy_tools/tree/master/tools/corals. The Galaxy CoralSNP *Queue Genotype Workflow* tool interface (B) consist of the analysis tools in the left tool panel. Selecting a tool displays the tool form in the center panel where the user can select the appropriate inputs for the tool and execute it. The tool outputs are added to the Galaxy analysis history on the right. The *Queue Genotype Workflow* tool accepts eight data files as inputs, the user metadata file and the Affymetrix sample attributes, annotation, summary, snp-posteriors, report, confidences and calls files. The tool includes a reference genome selection for the analysis. Once the tool is executed, the user can simply wait for the CoralSNP analysis pipeline to finish in the right panel.

The filtered VCF, population information file and user metadata were the inputs for the *Coral Multilocus Genotype* tool, which was executed through the R environment (RCoreTeam 2017). The VCF file was imported and converted into the genind format by the package vcfR v1.8.0 (Knaus and Grünwald 2017). The genind contains the individual genotypes that was then converted into a genclone format utilized by poppr v2.8.3 for clone identification (Kamvar et al. 2015; Kamvar et al. 2014). A distance matrix was calculated within poppr using the Prevosti’s absolute genetic distance (Prevosti et al. 1975), or the number of allelic differences between two individuals. From the distance matrix, known clone mates (ramets of the same genet) or replicate extractions from the same sample (Table S1) were compared to define a threshold for genet detection. This threshold encompasses technical (ie. missing alleles, genotyping error or DNA extraction differences) and biological (ie. somatic mutation) variation. The threshold was applied using mlg.fitler in poppr resulting in the assignment of samples to multi-locus genotype IDs, or genet IDs. Samples assigned to a genet ID with previously genotyped samples in the database took on the previous genet ID (ex. HG0000), whereas samples without matches to previously genotyped samples were assigned new genet IDs. The representative sample of the new genet ID was identified using the clonecorrect function in poppr.

The genetic distance matrix was used to calculate a neighbor-joining tree with 100 bootstrap replicates using the aboot function in poppr. An identity-by-state analysis was performed using SNPRelate as previously described (Kitchen et al. 2019; Zheng et al. 2012). The representative sample for each genet ID (*n*=193, excluding the genome samples, offspring of a Curacao cross with sample ID = SWSA, and plate 9SR22844), was used to identify populations with ADMIXTURE v1.3.0 (Alexander et al. 2009) outside of the Galaxy portal. Plate 9SR22844 was excluded due to higher percentage of missing data (average 1.271 ± 0.581% out of 96 samples, Figure S1E) and heterozygosity (average 14.163 ± 0.756 % in *A. palmata* and 12.875 ± 1.020 % in *A. cervicornis*, Figure S2A) for the entire plate that contained only Puerto Rico samples compared to the Puerto Rico samples on plate P9SR10076 (average missing data of 0.501 ± 0.251% out of 73 samples and average heterozygosity of 13.801 ± 0.626 % in *A. palmata* and 9.623 ± 0.446 in *A. cervicornis*, Figure S1C). The VCF file exported from Galaxy was filtered for representative genets and loci were reduced after applying a minor allele threshold of 0.05 with VCFtools, and converted using PLINK v1.9 (Chang et al. 2015). First, all representative genets were analyzed with inferred population of K=2 from 20 replicates with different random seeds to identify hybrids. Second, the two species were split from the filtered VCF. Populations of K ranging from 2 to 10 were run on each species separately over 20 replicates with different random seeds. In each iteration of ADMIXTURE, the replicates were combined and merged using the CLUMPAK server (Kopelman et al. 2015).

To determine the species of each sample, the genotypes were extracted from the VCF file using the extract.gt tool in the vcfR package. The nominally fixed probes were filtered further based on data from three plates where allele calls shared by less than 90% of the samples of a species were removed. Missing data was calculated for the entire probe set and the fixed probe set. The percentage of heterozygous alleles (AB) and the percentage of alleles matching each species in the fixed probe set was calculated from the reference (AA) or alternative (BB) alleles assigned for the *A. cervicornis* samples. A sample was identified as *A. palmata* or *A. cervicornis* if more than 85% of the fixed alleles match the respective species. Hybrid samples were identified as having 40% or greater heterozygosity.

A series of tables were generated from the analysis and imported into the respective database tables using the *Update STAG Database* tool (Figure 3A). This tool parses the metadata and genet information to append new records to the postgresql database (Figure S3).

### Galaxy CoralSNP Analysis Environment

The Galaxy Scientific Gateway called CoralSNP (https://coralsnp.science.psu.edu/galaxy) enables streamlined analysis of the Affymetrix genotype data described above to ultimately provide the user with a genet ID, converted raw genotype data, sample relatedness and hybrid status (Figure 1). A baseline set of reports (https://coralsnp.science.psu.edu/reports) provides various views of the data, and additional reports will be added over time.

The process is straightforward and shown in Figure 3B. A sample metadata file is created by the user using a template form (http://baumslab.org/research/data/). The metadata file contains a field where the user can choose when their data becomes publicly available, allowing a year hold. The user then uploads their raw Affymetrix data files into the Galaxy CoralSNP environment using the *Upload File* tool in the Galaxy tool panel. Next, the user selects the appropriate files as the inputs to the *Queue Genotype Workflow* tool (Figure 3B), which validates the metadata (*Validate Affy Metadata* tool), executes the CoralSNP workflow (Figure 3A) and updates a dataset that contains all previously genotyped samples as well as the STAG database (Figure S3) with the samples in the current run (*Update STAG Database* tool). From the user’s perspective, the entire analysis is as simple as uploading data and specifying it as the input to execute a tool.

The *Queue Genotype Workflow* tool shields the complexity of the analysis from the user and performs its functions via the Galaxy REST API. The CoralSNP workflow requires access to a dataset that contains all previously genotyped samples, so the tool imports this dataset into the user’s current Galaxy history from a Galaxy Data Library (https://coralsnp.science.psu.edu/galaxy/library/list#folders/Fcba2ba6d6fdc5d84). It is imperative that the previously genotyped samples contained within this VCF file are synchronized with the previously genotyped sample records contained within the STAG database. The *Ensure Synced* tool confirms that the data contained within these two components is synchronized before proceeding. The tool makes backup copies of the VCF file and the database before updating either component. Since both components are updated, multiple simultaneous analyses cannot be performed. The *Queue genotype workflow* tool handles this by ensuring that multiple simultaneous executions are handled serially. This is done by polling the status of the first execution until it has completed. Additional simultaneous executions are queued in the order in which they were submitted. If an analysis ends in an error with either the VCF file or the database updated so that they are no longer in sync, the backup copy of the appropriate component can be used to replace the problematic one in preparation for the next run.

The Galaxy CoralSNP environment contains an independent tool named *Export All Sample Data*, which produces a tabular dataset consisting of all samples and associated metadata in the STAG database. This dataset can be saved locally for analysis within other environments. The dataset that contains all previously genotyped samples can also be downloaded from the Galaxy Data Library, providing more options for additional analyses outside of Galaxy.

All the code and configuration files needed for hosting a local Galaxy CoralSNP instance are available in GitHub, and the instructions for configuring the environment are here https://github.com/gregvonkuster/galaxy_tools/blob/master/galaxy/README. The CoralSNP Science Gateway is hosted on a high-performance compute cluster environment managed by the Information Technology VM Hosting team at Pennsylvania State University.

### Symbiont genotyping: strain identification and background genera detection

The symbiont genotype data was analyzed in a similar manner to the coral data, but outside the Galaxy environment. Symbiont genotyping probes were identified from the BPW of all five plates after additional filtering to remove host contamination and low-resolution probes (*n*= 531, Table S6). The genotyping probes were subset from the full probe set using VCFtools and analyzed using a modified version of the *Coral Multilocus Genotype* tool. Notably, the ploidy was set to haploid. Because there was limited *a priori* information on the symbiont stain from microsatellite data, the distance threshold was set based on farthest and nearest threshold calculated by cutoff_predictor in poppr. Symbiont strains were given strain IDs in the format of SG0000.

For multiple vs. single strain detection from a single coral sample, five classification methods were used based on signal intensities of the filtered genotyping probes for samples assigned a strain ID. The intensities for each allele of each probe was extracted from the raw CEL file using Axiom Analysis Suite. Samples with prior symbiont genotyping from 12 to 13 microsatellites were used as the training set for all classification models where any sample with more than one allele per microsatellite marker was considered as containing multiple strains of *S. ‘fitti’* (*n*=17 samples with multiple strains and *n*=11 samples with a single strain). The remaining samples were the test set (*n*=265). The two data sets were centered and scaled prior to analysis.

The five classification tests included supervised learning models such as linear discriminant analysis (LDA) (MASS v7.3-51.4 R package, (Ripley 2002), decision tree (rpart v4.1-15 R package (Therneau and Atkinson 2019), and rpart.plot v3.0.8 R package (Milborrow 2019)), random forest (caret v6.0-84 R package, (Kuhn 2008)), and naïve Bayes (caret v6.0-84 R package, (Kuhn 2008)), and semi-supervised learning model using k nearest-neighbor masking 30% of the training data (SSC v2.0.0 R package, (González et al. 2019)). All tests, except for the LDA, were resampled three times with 10-fold cross-validation to evaluate model fit. The results of the five tests are presented as the percent of multiple strain assignment for each genotyped sample.

The background genera were assigned based on the fit of three of the classification tests above: LDA, decision tree and random forest. All samples and probes were first visualized in the Axiom Analysis Suite software to identify patterns in samples with known background symbiont populations (*A. cervicornis* with *Cladocopium: n= 2* (Lirman et al. 2014), *A. cervicornis* with *Durusdinium: n= 2*, Pacific acroporids with *Cladocopium: n*= 20 and *A. muricata* with *Durusdinium: n*= 5 (Hoadley et al. 2019)). Probes were filtered based on their recommended status (Table 1) and assignment of known samples above. A preliminary assignment of symbionts to genera was made for each sample based on those cluster patterns. The signal intensity for the genera probes (*n*= 18) was then extracted for all samples regardless of their genotype status using the Axiom Analysis Suite. The data was split into 80% for training and 20% for testing. Cross validation was performed on the decision tree and random forest models.

## Results

### Array Design and Validation

We identified 1.8 million high-quality coral SNPs that varied between the genomes of 42 previously sequenced *A. palmata* and *A. cervicornis* from four locations (Belize, Curacao, Florida, and U.S. Virgin Islands) using two variant callers, samtools mpileup (Li 2011) that uses likelihood scores and freebayes (Garrison and Marth 2012) that uses Bayesian posterior probabilities for variant calls. After Affymetrix filtered the 34,783 coral loci, the final array contained 32,124 loci with 53,579 probes, broken down into 25,889 fixed, 17,803 population and 9,887 *A. palmata* variable probes (Table 1 and Figure 2A). The majority of these variable sites are found within introns of coding sequences in the *A. digitifera* genome, followed by intergeneic regions (Figure 2B).

When comparing two deeply-sequenced *A. palmata* and *A. cervicornis* genomes to the reference *S. tridanidornium* genome, we identified 2,657 high-quality SNPs using samtools mpileup (Li 2011). When comparing 42 coral genome samples including the two above (Kitchen et al. 2019) to the draft genome of *A. cervicornis* ‘like’ *S. ‘fitti’*, 60,946 SNPs were considered high-quality (Reich et al, In Prep). Applying similar filtering methods to identify so-called ‘fixed’ differences between strains and populations as was done in the coral, we were left with only a small fraction of SNPs. Given the status of the *S. ‘fitti’* genome analysis at the time of the array design, we submitted more probes from the first comparison than the latter (2,269 from first comparison and 380 from the second comparison). Those loci were mostly found in the intergenic regions of the *Symbiodinium* genomes (Figure 2B). Of the 2,661 symbiont loci we submitted, all were retained in the final array with 4,021 probes covering fixed (*n*=3,663), population (*n*=304) and genera (*n*=54) categories (Table 1).

The recommended coral probes from the first plate were designated as the genotyping probes for the Caribbean acroporids in all subsequent analyses (Table 1). For the symbionts, all samples from the five plates that passed quality filtering (*n*=293 samples) were re-analyzed together using the BPW. The recommended probes were reduced further after removing probes that matched draft genome assemblies of *A. palmata*, *A. cervicornis* (Kitchen, unpublished), *A. tenuis* (Liew et al. 2016), *A. hyacinthus* (Liew et al. 2016), and *A. millepora* (Fuller et al. 2019) with high homology (blastn, e-value 1e-13), were not classified as Poly High Resolution, and had limited resolution outside of Florida samples (see Table S6). In particular, there were 146 probes that only distinguish the deeply-sequenced *A. cervicornis* symbiont strain, 247 probes that only distinguish the deeply-sequenced *A. palmata* symbiont strain, and 944 probes that distinguish the Florida *A. cervicornis* symbiont strains (*n*= 36 samples) from all the other samples. This resulted in 531 symbiont genotyping probes for downstream analysis.

The genotype success for each plate is presented in Table 2. The quality was first assessed by the background fluorescence of the non-polymorphic probes, or dish quality with a threshold of 82%. Then, only the samples with a call rate of 97% for the coral or symbiont probes, respectively, proceeded to the next step in the analysis. Because some of the samples were symbiont-enriched DNA or exclusively symbiont culture DNA, they failed BPW for the coral probe set. Alternatively, coral sperm and larvae failed the symbiont probe set (Table 2). Overall, Caribbean coral genotype calling was successful for samples with DNA concentrations as low as 0.064 ng/µl and as high 203.34 ng/µl (Table S1). Symbiont genotype calling worked for samples with DNA concentrations ranging from 0.23 to 203.34 ng/µl (Table S1).

### Coral genotyping via analysis portal

Four hundred seventy-nine corals (out of 520) were successfully genotyped using the genotyping probe set (Table 2 and Figure 4A) in the Galaxy CoralSNP analysis environment. The missing data ranged from 0.06% to 3.22% for the samples analyzed on the array (Figure S1). Plates differed in the amount of missing data that we attributed to a batch effect of sample preparation, but not sample preservative or extraction method because these were shared between plates. A significant positive correlation was detected between percent missing data and percent heterozygosity for each species (Pearson’s Correlation, *A. palmata* R^2^ = 0.4507, *p*= 8.142e-14; *A. cervicornis* R^2^ = 0.8223, *p* < 2.2e-16; Figure S2A), both of which are indications of sample quality. Misclassification of heterozygous calls can occur in samples with lower quality (Hong et al. 2008; Hong et al. 2012).

**Figure 4.**
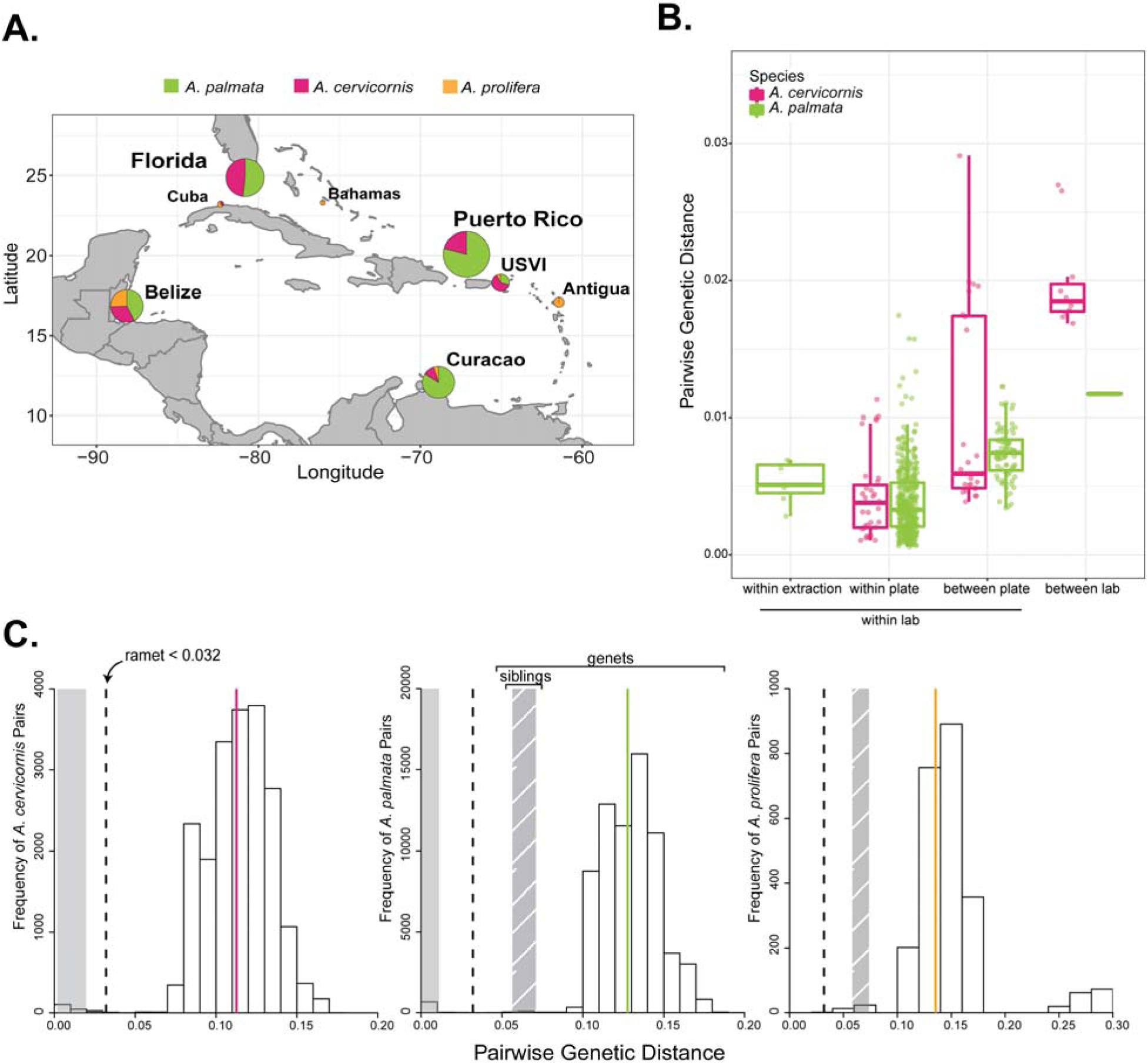
Caribbean acroporid genet identification. Pie-charts on the map of the Caribbean represent the percentage of species at each collection locations for the 479 genotyped samples (A). Prevosti’s pairwise genetic distance of ramets, or clone mates, was compared between technical replicates, samples within a plate and samples between plates processed within the same laboratory to those processed in a different laboratory (B). A histogram of the frequency of pairwise genetic distance values for each species indicates a break between ramets and genets (C). The dashed line is the threshold for ramet identification and the solid line is the average genetic distance for genets in the taxon (pink= *A.cervicornis*, green= *A. palmata*, orange= *A*. *prolifera*). The solid grey and hatch-marked grey shaded areas represent the mean ± standard deviation for ramets and siblings for each taxon, respectively.

Technical variation between replicate runs of the same DNA was low with an average genetic distance of 0.0053 ± 0.0015 between technical replicates (Mean ± 1 SD; samples SI-1, SI-10, SI-12, Table S8 and Figure 4B). The average pairwise genetic distance of ramets from the same genet (clone mates) within a plate was 0.0038 ± 0.0026 and between plates was 0.0079 ± 0.0041 (Figure 4B). Due to the larger genetic distances between technical replicates than ramets observed within a plate, we tested for differences in the five plates. There was a significant effect of plate on the genetic distance of ramets analyzed within plate (1-way ANOVA, F (4,391) = 17.58, *p*= 2.81e-13). Plate 9SR22843, which contained the technical replicates, had larger average pairwise genetic distances between ramets and technical replicates within the plate compared to three of the other plates (Tukey HSD, 9SR22843 was on average 0.0014 larger than 9SR22844 *p*=0.0003; 9SR22843 was on average 0.0015 larger than P9SR10073 p= 0.0019; 9SR22843 was on average 0.0025 larger than P9SR10076 *p*= 0.0000).

The threshold for genet assignment of samples was defined using previously identified ramets, ranging from two to six ramets per genet (shared baums_coral_genet_id in Table S1). The largest genetic distance within known ramets was ca. 0.0312 between a genome sample and array sample (ie. 14120_Mixed and 4960, Table S9). We used pairwise genetic distance = 0.032 as the threshold for genet assignment based on the observations above (Figure 4C). The average pairwise genetic distance among ramets was 0.0064 ± 0.0064 for all genet IDs with more than one ramet and ranged from 0.0006 to 0.0312 (Table S9). Additionally, tissue from eight genets extracted in two different laboratories recovered the same genet ID, albeit with differences in DNA concentration, missing data, and percent heterozygosity of the fixed probes (Figure 4B and Table S10). There was between 0.012 to 0.027 pairwise genetic distance among ramets of the same genet in this set, which is like what was observed for differences in genotyping methods (genome sequencing vs. array) and is within the genet threshold.

Between genet pairwise distance was on average 0.113 ± 0.023 for *A. cervicornis* and 0.128 ± 0.025 for *A. palmata* (Figure 4C). In the case of siblings from outcrossed offspring, the genetic distance ranged from 0.047 (SWSA-140 and SWSA-124) to 0.078 (SWSA-105 and SWSA-128) with an average genetic distance of 0.0642 ± 0.0068 (Figure 4C). Heterozygosity also varied by species and geographic region, ranging from 0.110 to 0.127 in *A. cervicornis* and 0.132 to 0.156 in *A. palmata* (Table 3 and Figure S2B). The inbreeding coefficient F_IS,_ which calculates the proportion of alleles within an individual that are shared with the population, was highest in Belize and Florida in both species (Table 3).

Genet resolution was reproducible across collection years, plates and different laboratories (Figure 4B, Figure 5 and Table S10). For example, HG0127 and HG0170 were recovered from samples collected between 2005 to 2018 and run on two different plates (Figure 5B). There was only one case where a genet defined via microsatellite genotyping was split into two genets as defined via SNP genotyping (blue lineage in Figure 5B). In the inverse situation, there were four cases where genets defined via microsatellite genotyping were no longer considered to be unique genets and combined with other samples defined via SNP genotyping (Table S1).

A Neighbor-Joining tree (Figure 5A) using the Prevosti’s genetic distance and identity-by-state analysis (Figure S4) clustered the samples, first by species and then by their collection location. However, the geographic regions were not clearly delineated using these methods. We could recover population clusters using an unsupervised model-based approach with ADMIXTURE (Figure 5C). After genet correction and applying a minor allele threshold of 5%, 18,823, 7,019, and 6,097 coral loci remain for all three taxa (n=193 samples), *A. palmata* (n= 90 samples) and *A. cervicornis* (n=64 samples), respectively The ancestry of each sample was assessed assuming two source populations for the full dataset and two to ten populations for each species separately. For K=2 of the entire dataset, the two species clearly separate with the hybrids having mixed ancestry (Figure 5C). The lowest prediction error for *A. cervicornis* was three inferred populations (Figure S5) with a population in Florida, a population in Belize and a population in USVI and Puerto Rico (Figure 5C). Three populations were also predicted in *A. palmata* with a population in Florida and Belize, a population in Puerto Rico and a population in the Curacao (Figures S5 and 5C).

**Figure 5.**
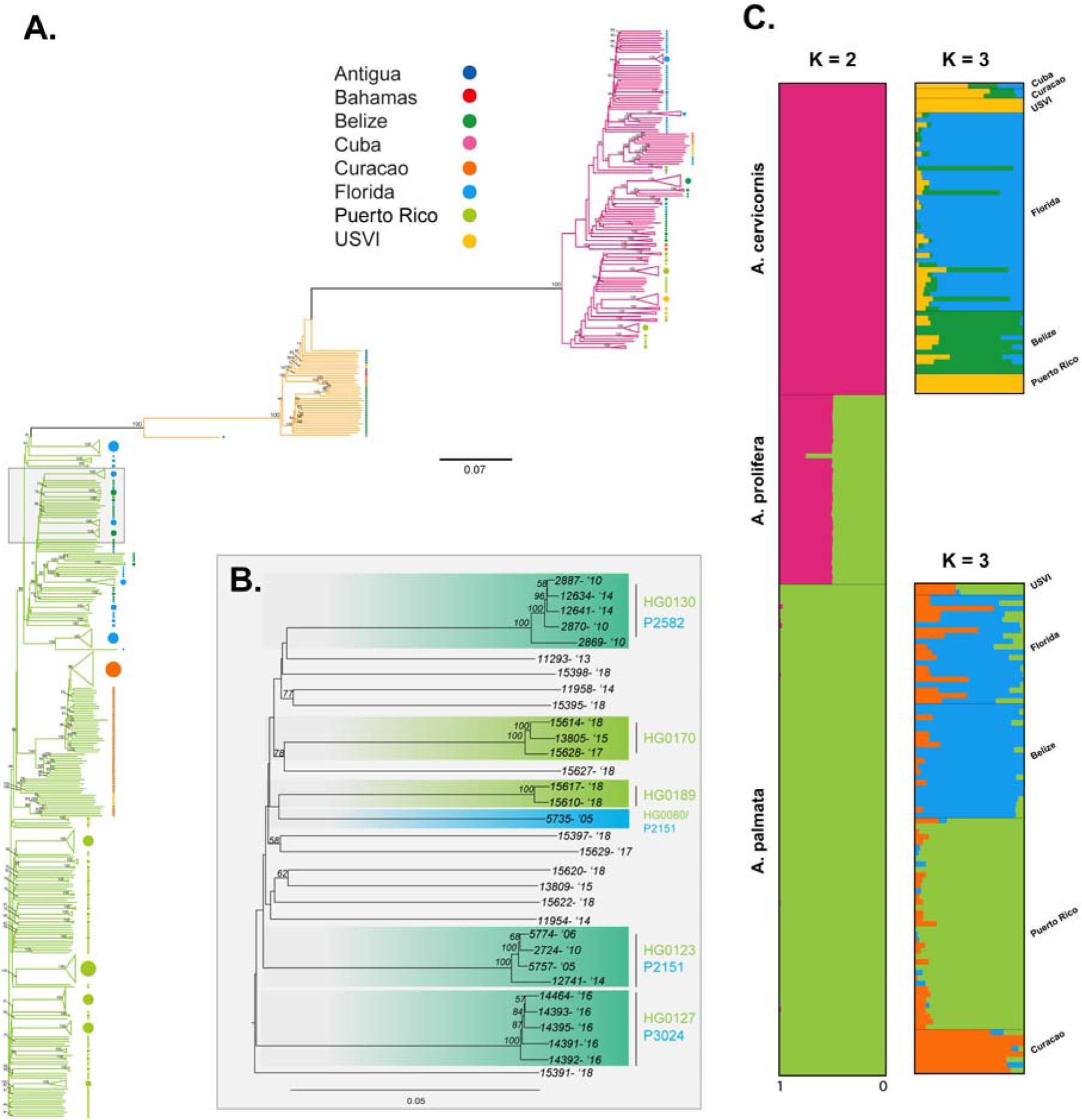
Caribbean acroporid population analysis. Prevosti’s genetic distance of 19,694 SNPs was used to construct a neighbor-joining tree (A). The branches are colored by their genetic species identification and collection locations are indicated by the color of the circle at the terminal ends (Antigua = blue, Bahamas = red, Belize =green, Cuba = pink, Curacao = orange, Florida = light blue, Puerto Rico = light green, and USVI= yellow). Nodal support is based on the 100 bootstrap replicates. The nodes of genets with multiple ramets identified with the SNP data are collapsed in the tree. In panel B, an example of genet resolution is provided based on the array SNP data and the previous microsatellite IDs over different collection years. The SNP genet ID is presented in green on the top and the microsatellite genet ID is presented in blue on the bottom. The clades are shaded blue-green where the two genotyping methods are congruent. The collection year is presented next to the sample identification number. ADMIXTURE was run on a representative sample for each genet (*n*=193), excluding genome samples, offspring of a Curacao cross and Puerto Rico samples from plate 9SR22844 (C). Individual bars represent the relative proportion of membership of a sample to the inferred K populations. Results from two source populations for all samples and three source populations for each species separately (K=3 had the lowest cross-validation error for both species, Figure S5).

### Hybrid identification

The genetic species assignment was based on 9,072 fixed probes. The proportion of ancestry from each parental species was calculated for each sample and used to identify hybrids (Figure 6). There were 39 *A. prolifera* hybrids of which all but one appears to be a F1 hybrid (Figure 5C and Figure 6). Based on the field calls, one hybrid detected with the array data was previously misidentified as *A. palmata* and 11 samples identified as hybrids in the field (*n*=7 larvae and *n*=4 adults) were assigned to one of the parental species instead.

**Figure 6.**
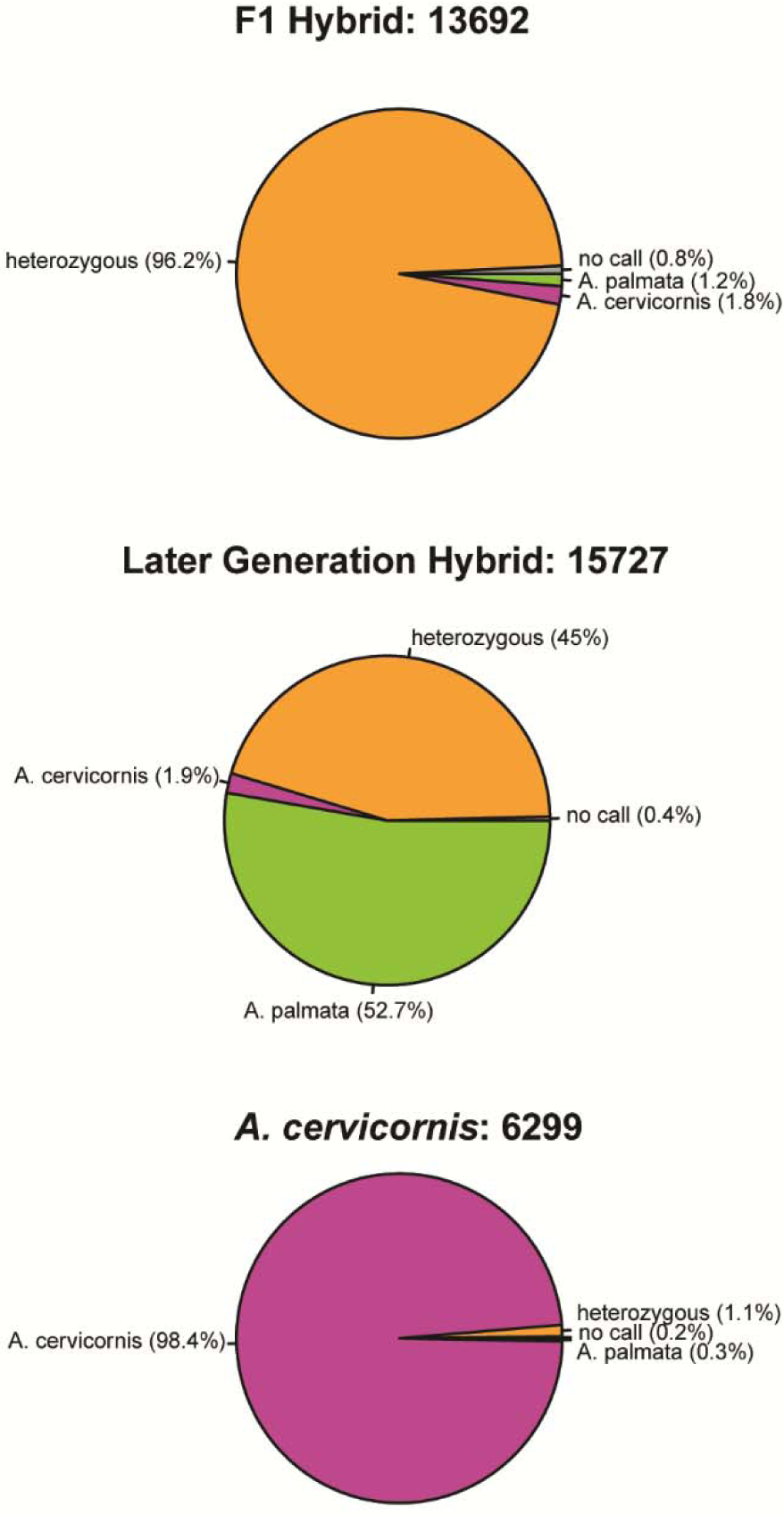
Species-specific SNPs identify hybrids. Sample 13692 was identified as an F1 and sample 15727 as a later generation hybrid. For comparison, sample 6299 is identified as a pure *A. cervicornis* sample. The 9,072 fixed SNPs were scored as homozygous for each species, *A. palmata* or *A. cervicornis*, or as heterozygous.

### Symbiont Genotyping

There were 293 samples that passed the BPW for the symbiont probes. Unlike the coral samples, the extraction method mattered for symbiont DNA recovery and genotyping. This is exemplified by the failure of all but one replicate DNA extractions using the magnetic bead protocol and successful genotyping of all samples after DNA extraction with the QIAGEN DNeasy kit. 186 putative *S. ‘fitti’* strains were identified based on a genetic distance threshold of 0.0018. We call these putative strains based on the limited *a priori* information available for setting the strain detection threshold. Enriched symbiont DNA and mixed DNA extractions from the same tissue shared the same strain ID as did technical replicates of the same DNA extractions from the same ramet (Table S1).

Sometimes more than one strain can be present in a given host and the strain ID might represent a mixture of different *S. ‘fitti’* strains. We attempted to identify colonization of single or multiple strains in a host sample through various supervised and semi-supervised classification methods using the signal intensity of the symbiont genotyping probes. The posterior probabilities of the LDA were used to determine likely colonization status for the known and unknown samples (Figure S6A). There was a difference in the distribution of the multiple and single colonized samples on LD1 (Figure S6A); however, two single strain samples overlapped the distribution of samples with multiple strains. More unknown samples overlapped with the distribution of samples with multiple strains compared to the distribution of samples with a single strain (Figure S6A). The decision tree had an accuracy of 53.6% and only required signal intensity of two probes for the classification with the lowest cross-validation error (probes AX.197983721.B and AX.198082605.A, Figure S6B). For the random forest model, the accuracy was estimated to be 66.9% with higher classification error for the single strain samples (multiple error = 29.4%, single error = 54.5%). Five trees were predicted to have the lowest error with the largest number of nodes, one of which is presented in Figure S6C. Naïve Bayes had an accuracy of 69.2% for the training data. Lastly, the semi-supervised k-nearest neighbor model had an accuracy of 65.8%. The results of all classification models were calculated as the percent agreement of multiple strains prediction (ex. 2 out of 5 tests predicted multiple strains = 40%).

There were 112 samples that were likely colonized by a single strain (0-20% agreement for multiple) and 157 samples that were likely colonized by multiple strains (80-100% agreement for multiple) (Table S1).

In addition to multiple strains of *S. ‘fitti’* present in a single coral host, the coral can be colonized by additional symbiont genera. We used the same classification methods above to detect background genera using the signal intensity of 18 genera probes (Table S4), but each sample was pre-assigned to a genus or classified as not colonized based on their allele patterns. The prediction accuracy of the LDA (Figure 7), decision tree (Figure S7A) and random forest (Figure S7B) was 98.9%, 96.4% and 98.9%, respectively. The predictions for each model are presented in Table S1. The presence of *Breviolum* was detected in thirteen samples with one of the classification methods, ranging from 0.2% to 100% probability. Of these, seven had probabilities greater than 60% and two of those also had *S. ‘fitti’* strain IDs indicating co-infection. The *Cladocopium* containing samples were split into two clusters, one contained samples that were exclusively *A. muricata* hosts (Cladocopium 2) and the other contained host samples that were *A. cervicornis* (*n*= 2), *A. digitifera* (*n*= 8), and *A. millepora* (*n*= 5). Finally, there were 49 samples with *Durisdinium* (*n*= 5 *A. muricata*, 3 *A.cervicornis*, 41 *A. palmata*). Samples containing *Cladocopium* or *Durisdinium* failed the *S. ‘fitti’* genotyping analysis.

**Figure 7.**
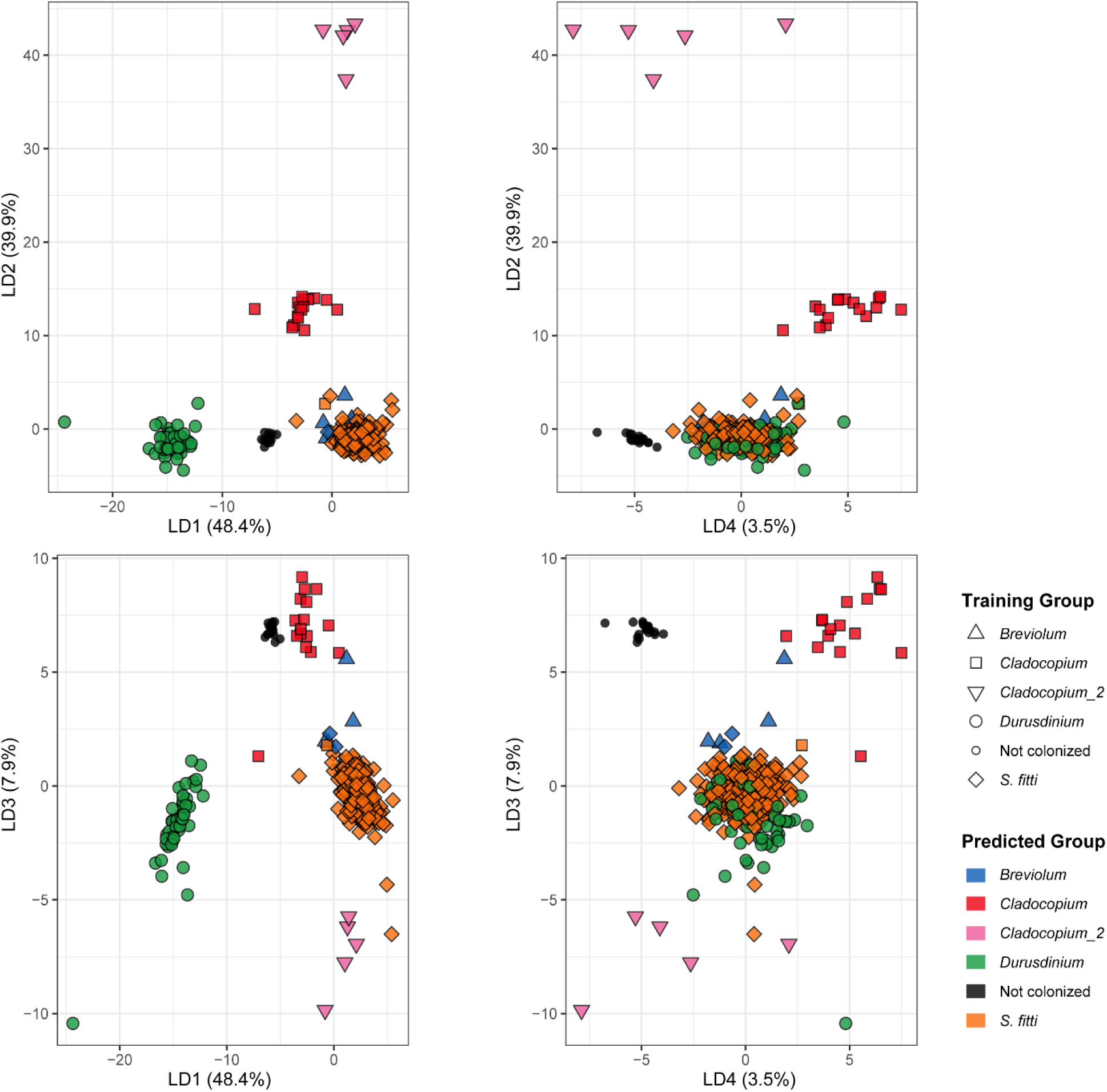
Detection of background symbiont genera. Results of the linear discriminant analysis where the shape denotes the preliminary training group genera assignment and the color is the predicted genera assignment. LD1 separates *Durusdinium* and not colonized samples, LD2 separates *Cladocopium* taxa and LD3 starts to separate *Breviolum* from the *Symbiodinium* group.

### Suitability for Pacific acroporids

Based on *in silico* genome searches, 26,963 of the coral probes matched *A. hyacinthus*, 28,395 matched *A. millepora* and 14,399 matched *A. tenuis.* Given that our probes were designed using the genome assembly of *A. digitifera* and that they had high homology to other species, we tested whether we could find a conserved set of probes across the Pacific acroproids for future genotyping studies. The Pacific samples were run separately for each species in the genotyping mode in the Axiom Analysis Suite to get the recommended probe set for each species. This analysis did not enforce a dish-quality threshold. A total of 15,717, 21,520 and 7,275 probes were recommended for *A. digitifera* (*n*= 9 samples), *A. millepora* (*n*= 5 samples) and *A. muricata* (*n*= 11 samples), respectively. Only those probes that were recommended for all three species were used for further analysis (*n*= 1,779 probes, Table S7). The pairwise genetic distance among *A. digitifera* samples ranged from 0.018 to 0.081 (Figure 8A), with tight clustering in all but one sample. Two *A. millepora* samples were nearly identical (Prevosti’s distance = 0.00084) and differed only at two probes (Figure 8B), while the largest pairwise genetic distance was only 0.024 (difference of 42 probes). Similarly, two *A. muricata* samples were also closely related, with a Prevosti’s distance of 0.004 (Figure 8C). For this species a clear pattern emerged separating the nearshore and offshore samples with a maximum pairwise distance of 0.429 (763 probes, Figure 8C). Although the sample size is too limited for each species to determine genotyping thresholds, less than 50 loci are necessary to identify the 33 unique genets in this dataset based on a genotype accumulation curve (Figure S8).

**Figure 8.**
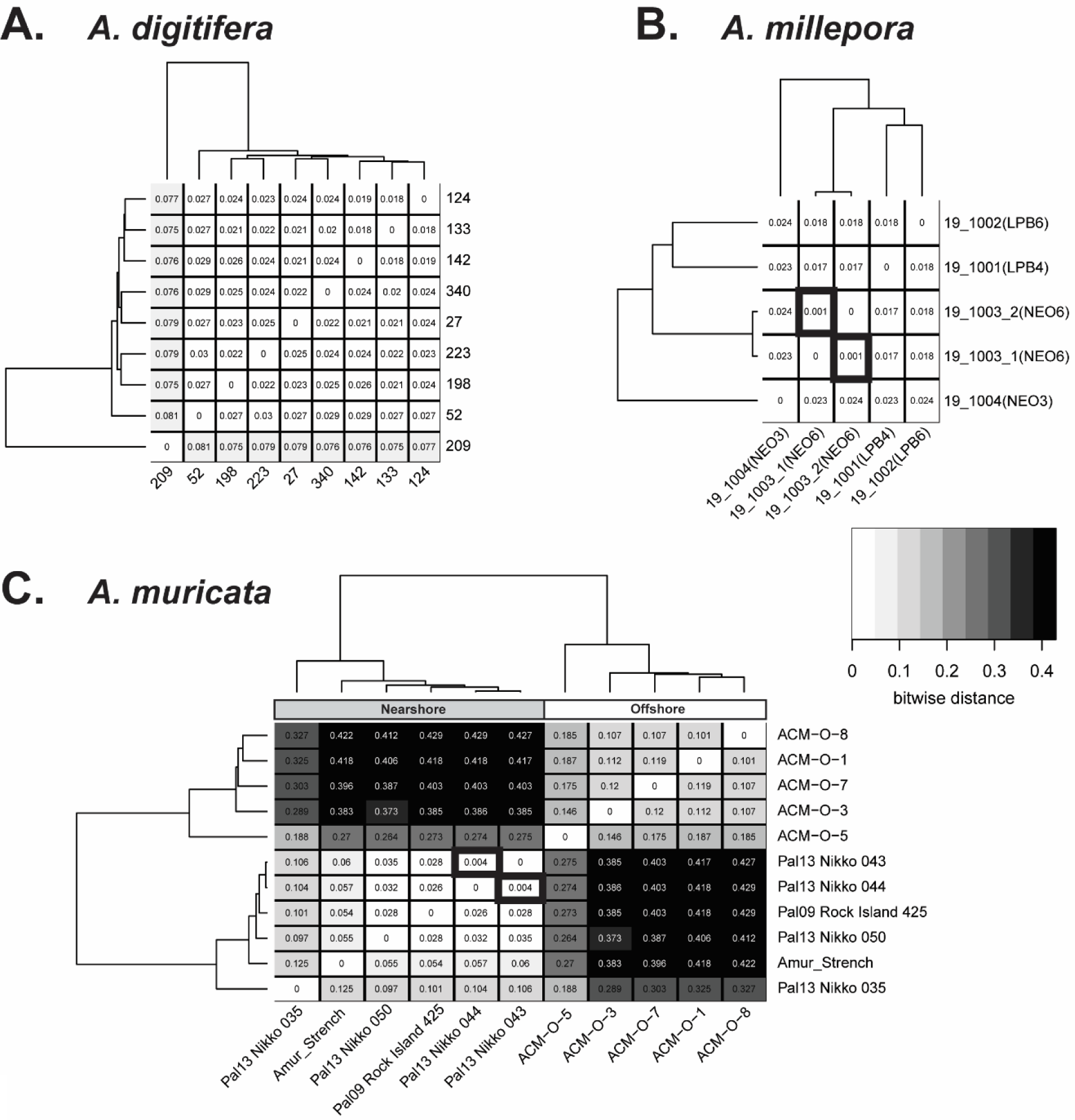
Genetic distance of Pacific acroporids using 1,779 shared probes. The relatedness of samples from three Pacific species, *A. digitifera* (A), *A. millepora* (B) and *A. muricata* (C) were compared using Prevosti’s genetic distance. The distance for each pairwise sample combination is displayed in the respective square of the heatmap. The darker the shading, the larger the genetic distance between samples. The dendrogram on the top and side represents the hierarchical clustering of the samples based on genetic relatedness. Samples with thick black borders are nearly identical for the probes tested and are likely the same genet. In the case of *A. muricata* (C), clear separation is observed of nearshore and offshore samples.

## Discussion

Here we report the first genotyping array for corals, which in combination with an open access Galaxy Scientific Gateway to execute the Standard Tools for Acroporid Genotyping (STAG) workflow produces multi-locus genotypes for coral hosts and their algal symbionts. In the workflow, new user-supplied samples are compared to previously genotyped samples and their results contribute to the growing STAG database (Figure 1). This archive of coral genets and symbiont strains can be used to identify reefs with high host and/or symbiont genetic diversity, temporal and spatial changes, and shuffling in host-symbiont pairings. In addition, a subset of the Caribbean genotyping probes can be used to genotype Pacific acroporids, expanding the utility of the STAG workflow to hundreds of species.

The SNP array and analysis workflow developed here delineate genets in agreement with the previous gold standard for Caribbean acroporid genotyping, multiplex microsatellite genotyping (Baums et al. 2005a). The STAG workflow uses 61% of the coral loci to produce the host genotype (Table 1) and identified 325 genets out of 479 genotyped samples (Table 3). The average genetic distance of 0.0064 (difference of 0.64%) among ramets was well below our maximum between genet genetic distance threshold of 0.032 (Figure 4C), which accounts for both biological processes (mutations) and technical error during genotyping. We estimate that technical error accounts for ≤ 0.0053 (0.53%) of this variation based on the lower genetic distance observed within plate for both species than the replicate analysis on the same DNA extraction from a single tissue sample (Figure 4B and Table S8). The differences observed in ramet genetic distance between plates may be due to the genotyping probe set applied to all plates irrespective of the recommended set for each plate (Table S5). Differences in genetic distances of ramets can also arise from DNA quality that is influenced by sample preservation, tissue type, extraction method, and extraction laboratory. We found a positive relationship between missing data and total heterozygosity (Figure S2), suggesting that a portion of heterozygous genotype calls in the lower quality samples might be an artifact of technical error. This was evident in the different percent heterozygous estimate of the fixed probes in the between laboratory replicate extractions (Table S10). However, our technical error is similar to previous genotype concordance estimates ranging from 0.2% to 2.4% for replicates of a given subject genotyped on Affymetrix SNP arrays for humans (Hong et al. 2012), rainbow trout (Palti et al. 2015), soybean (Lee et al. 2015) and walnut (Marrano et al. 2019). In that latter study, the variation was also higher between technical replicates than biological replicates, which the authors attributed to DNA quality. All these sources of technical variation are accounted for in the genotype assignment by the STAG workflow, resulting in robust coral genet identification.

Technical variability can be minimized by standardizing procedures. We recommend that adult samples of at least 3-4 polyps are preserved in 95% non-denatured ethanol (190 proof), stored as cold as possible and extracted using the Qiagen DNeasy tissue extraction kit. DNA requirements are modest for the Axiom SNP array. Adult tissue, single larva and concentrated sperm were successfully genotyped in samples with DNA concentrations as low as 63 pg/µl, although higher concentrations are recommended. While high-quality, non-degraded DNA provided the best results, moderately degraded samples (i.e extractions that show a dense band of high molecular weight DNA with some smearing across size ranges) were also successfully genotyped. DNA requirements with respect to quality and quantity are thus comparable to RADseq and whole genome sequencing techniques.

*A. palmata* and *A. cervicornis* differ in the scale of dispersal with *A. cervicornis* showing higher levels of population subdivision across the Caribbean and North Atlantic compared to *A. palmata* (Baums et al. 2014; Baums et al. 2010; Hemond and Vollmer 2010; Vollmer and Palumbi 2007; Drury et al. 2016; Porto-Hannes et al. 2014). *A. palmata* stands were found to be structured into two long-separated East/West populations based on microsatellite data (Baums et al. 2005b), but additional samples from the Mesoamerican Reef Tract (Porto-Hannes et al. 2014) and the development of SNP markers (Devlin-Durante and Baums 2017) resulted in the discovery of further population structure. Our results from a limited number of geographic locations identified three populations in *A. palmata* consistent with the previous study by Devlin-Durante and Baums (2017), recovering the East/West divide with additional substructure between Puerto Rico and Curacao in the East. We also recovered three populations in *A. cervicornis*, but with substructure detected between the Western Caribbean populations of Florida and Belize.

Quantifying the extent to which introgression has historically occurred and may occur now can elucidate the evolutionary and ecological significance of hybridization in acroporids. Using the species-specific fixed SNPs, we identified 39 F1 hybrid genets and corrected several species misidentifications in the field based on colony morphology (one classified hybrid identified as *A. palmata* in the field and two classified *A. palmata* identified as hybrid in the field). While F1 hybrids are more common, later generation backcrosses do occur (van Oppen et al. 2000; Vollmer and Palumbi 2002a) albeit the direction of introgression has been debated (Palumbi et al. 2012; Vollmer and Palumbi 2002b; 2003). Here, we identified one later generation hybrid that was classified as a putative backcross *A. palmata* (44.98% heterozygous and 52.7% *A. palmata;* Figure 6) in contrast to earlier findings that backcrosses are restricted to introgression of *A. palmata* genes into the *A. cervicornis* genome. A recent report also found putative *A. palmata* backcrosses based on microsatellite data in the Lesser Antilles (Japaud et al. 2019). Together, these results support the conclusion of bidirectional introgression in Caribbean acroporids.

Because of the intimate association between corals and algae, the SNP array was designed to assay host and symbiont DNA simultaneously, a novel application for the Axiom SNP array. The array contains a much smaller number of symbiont-specific probes compared to host probes and thus information gleaned from these probes is more limited. The large genome size, haploidy and asexuality of *Symbiodinium ‘fitti’*, the dominant symbiont of the Caribbean acroporids (Pinzón et al. 2011), presents challenges. The lower allelic diversity of *S. ‘fitti’* microsatellite loci compared to the allele diversity of their cnidarian host counterparts necessitates using larger number of loci for strain resolution (Baums et al. 2014). After exhaustive filtering of the symbiont genotyping probes based on their performance, only 20% of the loci remained which recovered reproducible strain identity in replicate ramets of a given genet. However, given the limited prior strain information for the samples, the conservative threshold we used for strain assignment will need to be validated with more known strains in the future. Only 58% of coral samples with symbionts yielded an *S. ‘fitti’* genotype (Table 2).

Failures were either due to inefficient symbiont DNA recovery in the extraction or to presence of other Symbiodiniaceae genera. Comparison of strain resolution achieved with the SNP array relative to microsatellite strain resolution revealed previously unresolved strain diversity. It is not yet clear how much of this strain diversity results from mutational processes versus diversity produced as a result of recombination between strains (Baums et al. 2014; Liu et al. 2018).

*Acropora* colonies are at times colonized by more than one strain of *S. ‘fitti’* (Baums et al. 2014) but classification of colonies as being colonized by a single or multiple strains was challenging (Fig S5). In contrast, the ability to detect the presence of other Symbiodiniaceae genera within coral samples is encouraging (Fig 7). We detected eight *A. cervicornis* and 44 *A. palmata* colonies that harbored symbionts of the genera *Breviolum*, *Cladocopium* or *Durusdinium*. Of these, three *A. cervicornis* and three *A. palmata* are likely to be co-colonized by *Breviolum* and *S. ‘fitti’*, a combination of symbionts shown to be intermittent in *A. cervicornis* through profiling the *ITS2* gene (Thornhill et al. 2006). Further, symbiont genera detected in nearshore (=*Durusdinium*) and offshore (=*Cladocopium*) *A. muricata* samples were consistent with a recent study by Hoadley et al. (2019), although this taxon of *Cladocopium* (Cladocopium_2) was distinctly different from the other *Cladocopium* taxon (Cladocopium) containing both Caribbean and Pacific hosts (Fig. 6). The two *Cladocopium* groups differed in their signal intensities for the genera probes with samples in the Cladocopium_2 having signal intensity on average 4.5x higher than samples within the Cladocopium taxon. Signal intensities may vary due to quantity of DNA, random difference in hybridization efficiency, and variable affinity of probes to different symbiont taxa within genera. Thus, we stress here that the SNP array cannot be used to derive quantitative differences among symbiont taxa associated with a coral sample. Moreover, DNA from cultured *S. tridacnidorium* was also on average 4x higher than mixed *Acropora*-*S. ‘fitti’* samples, suggesting that “pure” symbiont DNA extracts cannot be directly compared to mixed host-symbiont samples. Further experiments should benchmark the method by testing mixtures of Symbiodiniaceae genera with known composition.

Application of the current array to non-target Pacific acroporid species is possible when the sole intent is to delineate genets as is often required in restoration settings. Because of the large ascertainment bias inherent in applying probes designed for Caribbean acroporids to long-separated Pacific species, population genetic models and models designed to detect loci under selection should not be applied to this data.

The combination of the tools presented here provides reliable, standardized identification of host genotypes in diverse *Acropora* spp. and symbiont strains of the Caribbean species. These markers and analysis tools can be used for basic research questions such as gene by environment interactions, hybridization history, or identification of loci under selection. Genetic linkage maps can be generated and inbreeding levels, and relatedness questions can be addressed. Because of the low error rate, the SNP array is particularly suited for the detection of somatic mutation, which are expected to be common in the large, old genets that are now dominating Caribbean *Acropora* populations. Restoration practitioners can use the information to design propagule transfer zones and choose genets for nursery rearing.

## Acknowledgements

Funding for this project was supported by NOAA Office for Coastal Management NA17NOS4820083 awarded to IBB and SAK, and NSF OCE-1537959 awarded to IBB, NDF and WM. We are grateful for assistance with sample collection by Zachary Fuller, Mikhail Matz, Stephen Palumbi, Todd LaJeunesse, Jose M. Eirin-Lopez, Katie Flynn, Sea Ventures, Coral Restoration Foundation, The Nature Conservancy, and Fragments of Hope, and sample preparation by Meghann Devlin-Durante and Sam Piorkowski. We thank Sam Vohsen and Eslam Osman for valuable feedback on the Multilocus Genotype tool and Reef Futures 2018 workshop participants for their feedback on the Galaxy CoralSNP analysis environment. Permits for samples include Florida: CRF permit numbers CRF-2017-009, CRF-2017-012, NOAA FKNMS permit numbers FKNMS-2011-159-A4, FKNMS-2001-009, FKNMS-2014-148-A2, and FKNMS-2010-130-A, Belize: CITES Permit 0385, 7487 and 7488; Curacao: CITES Permit 16US784243/9 and 12US784243/9; Puerto Rico Department of Natural and Environmental Resources permit 2018-000064;and USVI Department of planning and natural resources, Division of fish and wildlife DFW14017T.

## Data Availability

The Galaxy CoralSNP analysis environment is available at and database reports are available at https://coralsnp.science.psu.edu/reports. The code for the new tools developed for this study are available at https://github.com/gregvonkuster/galaxy_tools/tree/master/tools/corals and https://github.com/gregvonkuster/galaxy_tools/tree/master/galaxy. Sequences for the genome samples are available on NCBI under SRA project SRP149363. The coral probe annotation is provided in Supplemental File 1 and the symbiont probe annotation is provided in Supplemental File 2.

## Author Contributions

SAK conceived the project, obtained funding, designed and validated the array, developed genotyping analysis tools and wrote the manuscript. GVK developed genotyping analysis tools, created the STAG database, built the custom Galaxy Science Gateway and wrote part of the manuscript. KVK organized and extracted samples and edited the manuscript. HGR identified the *S. ‘fitti’* SNPs and edited the manuscript. WM identified the *S. tridacnidorium* SNPs and obtained funding. SG provided Puerto Rico samples for analysis. NDF supplied coral samples, conducted laboratory comparison methods, obtained funding and edited the manuscript. IBB conceived the project, obtained funding, contributed to the array and database design and wrote the manuscript.

## Supplemental Material

**Figure S1.**
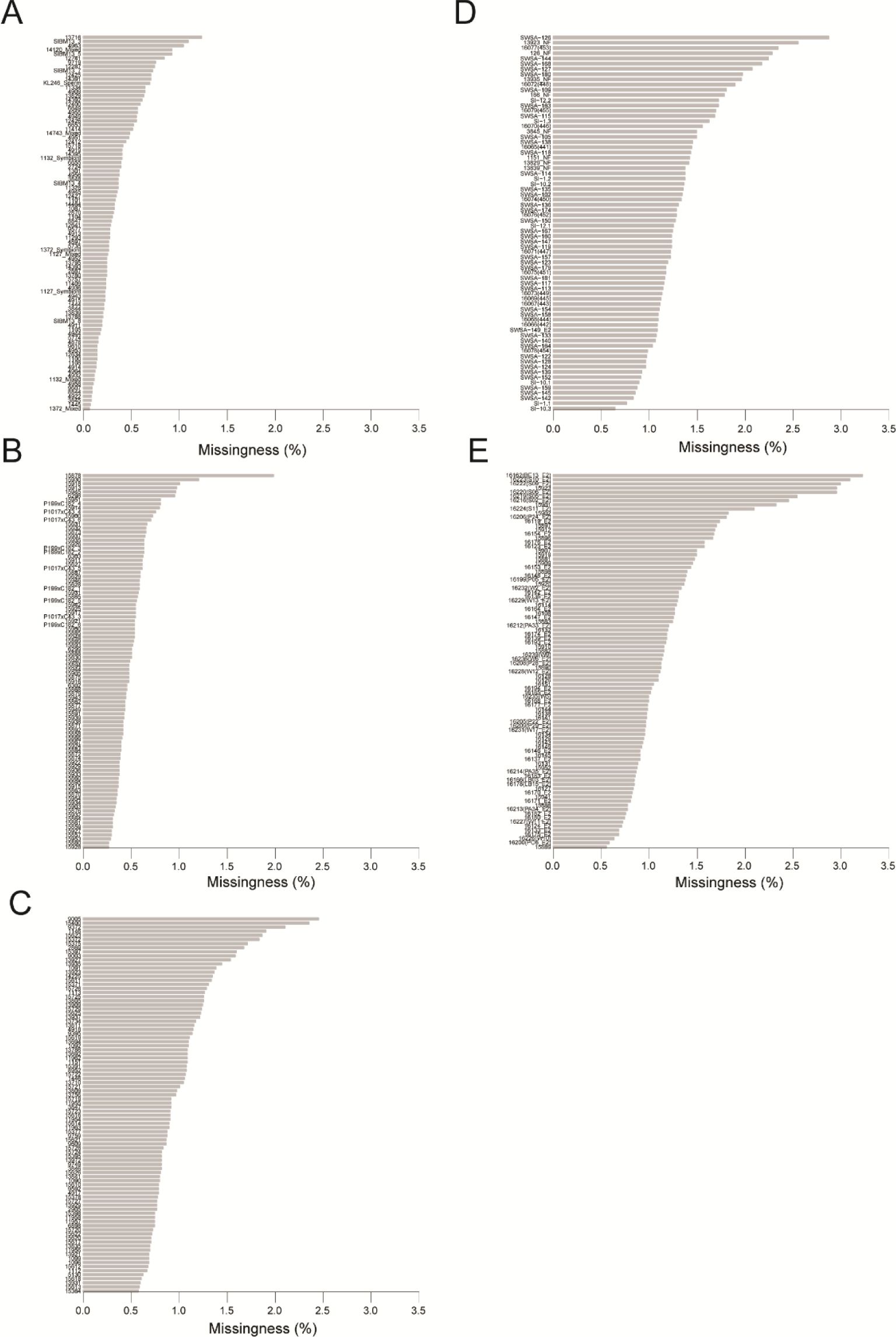
Percentage of missing genotype calls per sample split by each plate. Plates P9SR10073 (A), P9SR10074 (B), P9SR10076 (C), 9SR22843 (D) and 9SR22844 (E).

**Figure S2.**
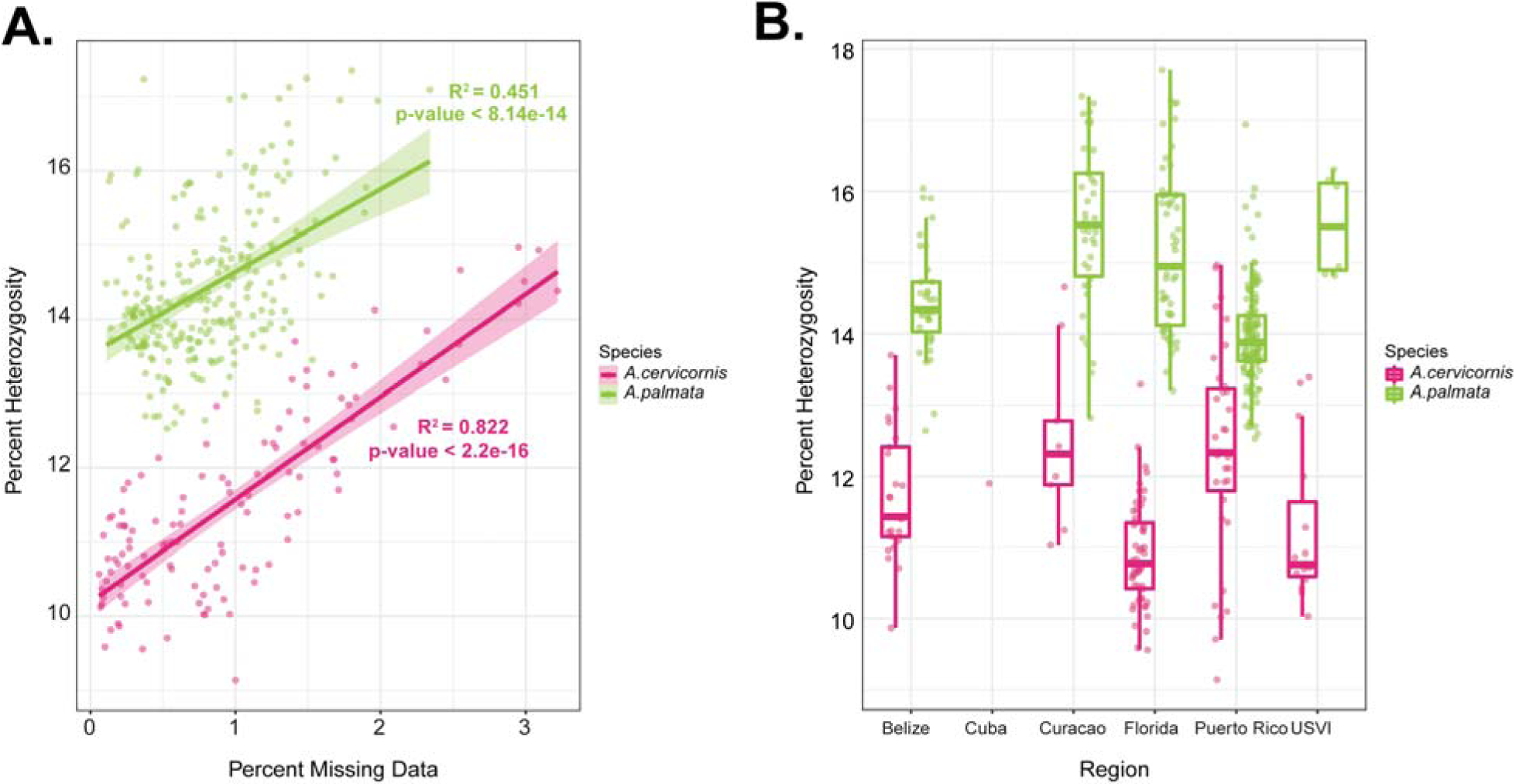
Percentage of heterozygosity by species and geographic region. A positive correlation was detected between percentage of missing data and heterozygosity for each species (A). A breakdown by collection location and species reveals higher total percent heterozygosity in *A. palmata* compared to *A. cervicornis* (B).

**Figure S3.**
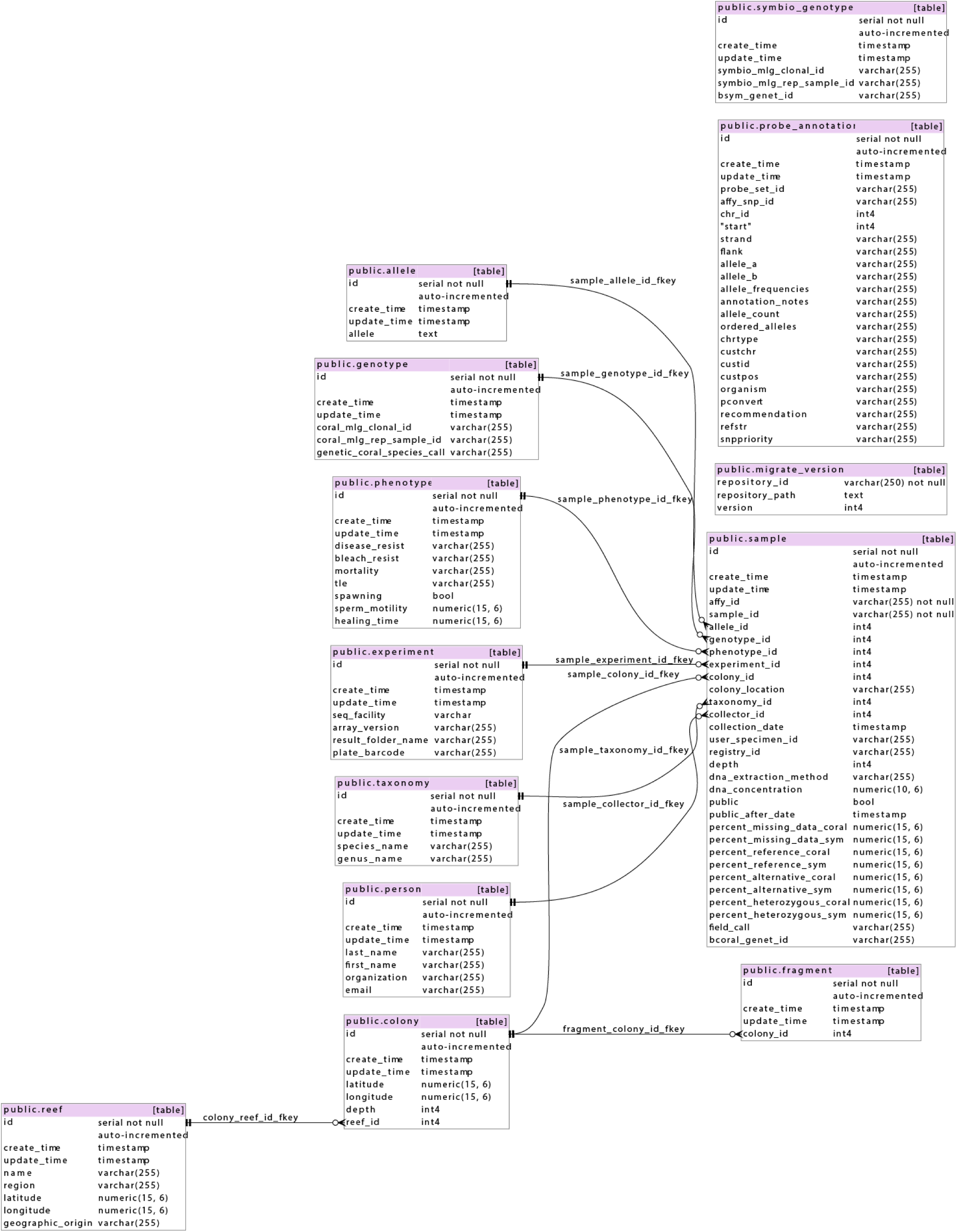
STAG database schema. This database was initially populated with the genotypes of 42 acroporid genomes that were sequenced in 2017. The database contains the genotype pattern for each unique clonal ID and a list of all samples matching that clonal ID. It also contains metadata provided by the user about each sample such as collection site (GPS), collection date, sample depth, contact information of the collector, and sequencing facility of the raw data. The Python code that creates the stag database is available on GitHub at https://github.com/gregvonkuster/galaxy_tools/blob/master/galaxy/corals_database/lib/galaxy/model/corals/mapping.py.

**Figure S4.**
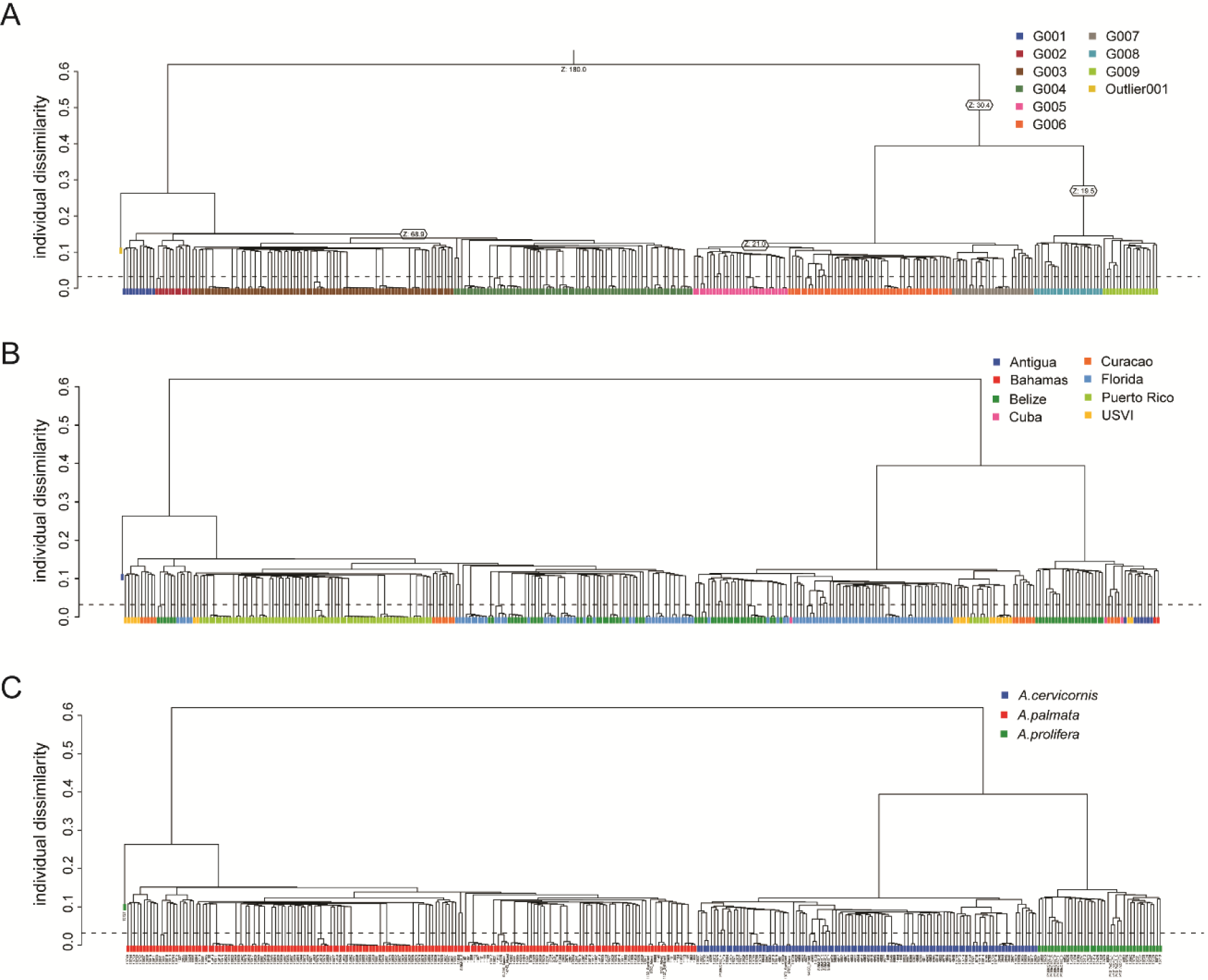
Identity-by-state clustering for three plates based on z-score (A), region (B) or species (C).

**Figure S5.**
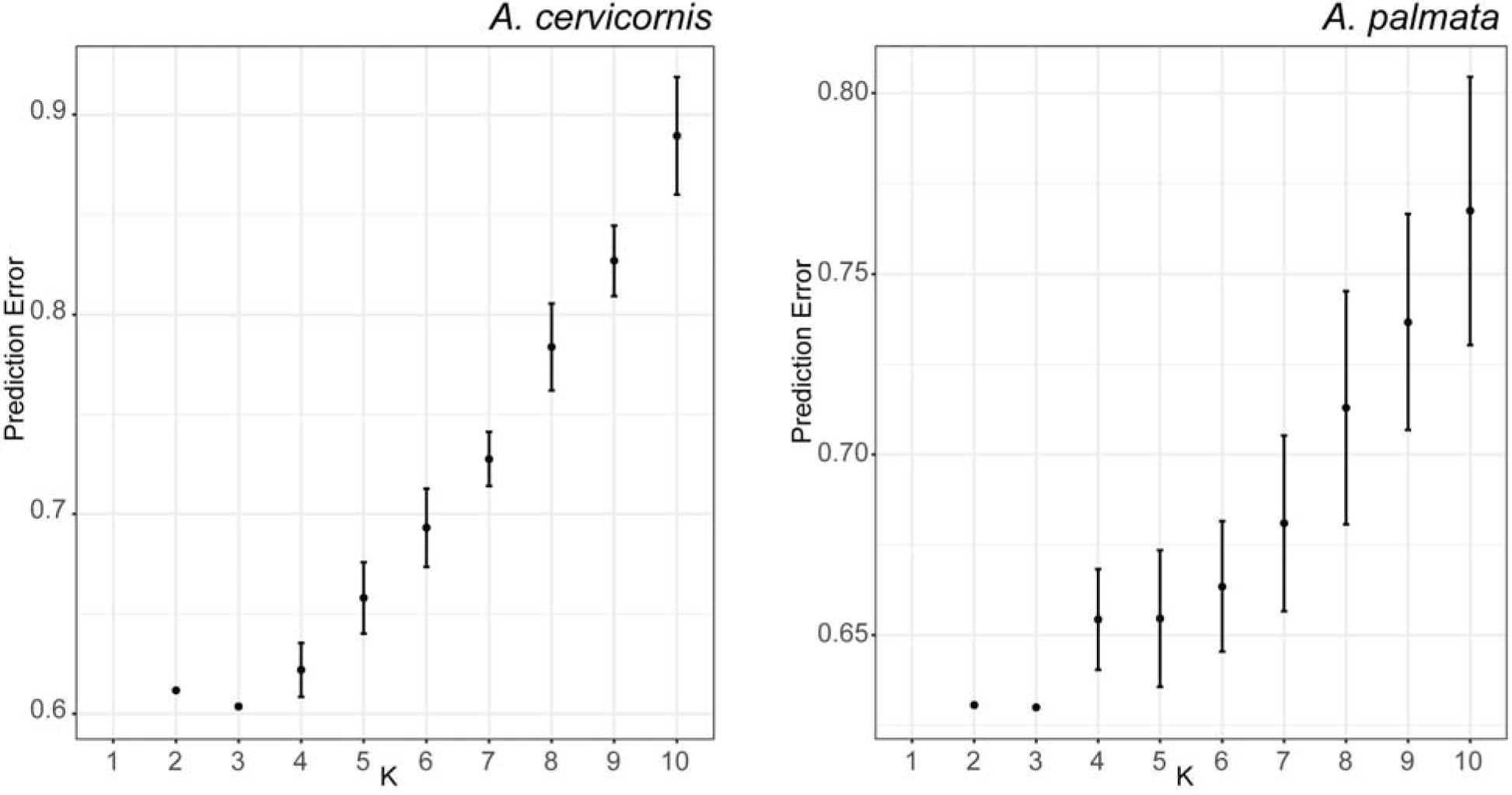
Cross-validation error of tested K populations. Each value of K was repeated 20 times with a different random seed in ADMIXTURE. The mean value of CV prediction error +/- the standard deviation is shown. K=3 had the lowest CV errors for both species (*A. cervicornis* = 0.604 ± 0.0009 and *A. palmata* 0.630 ± 0.0002).

**Figure S6.**
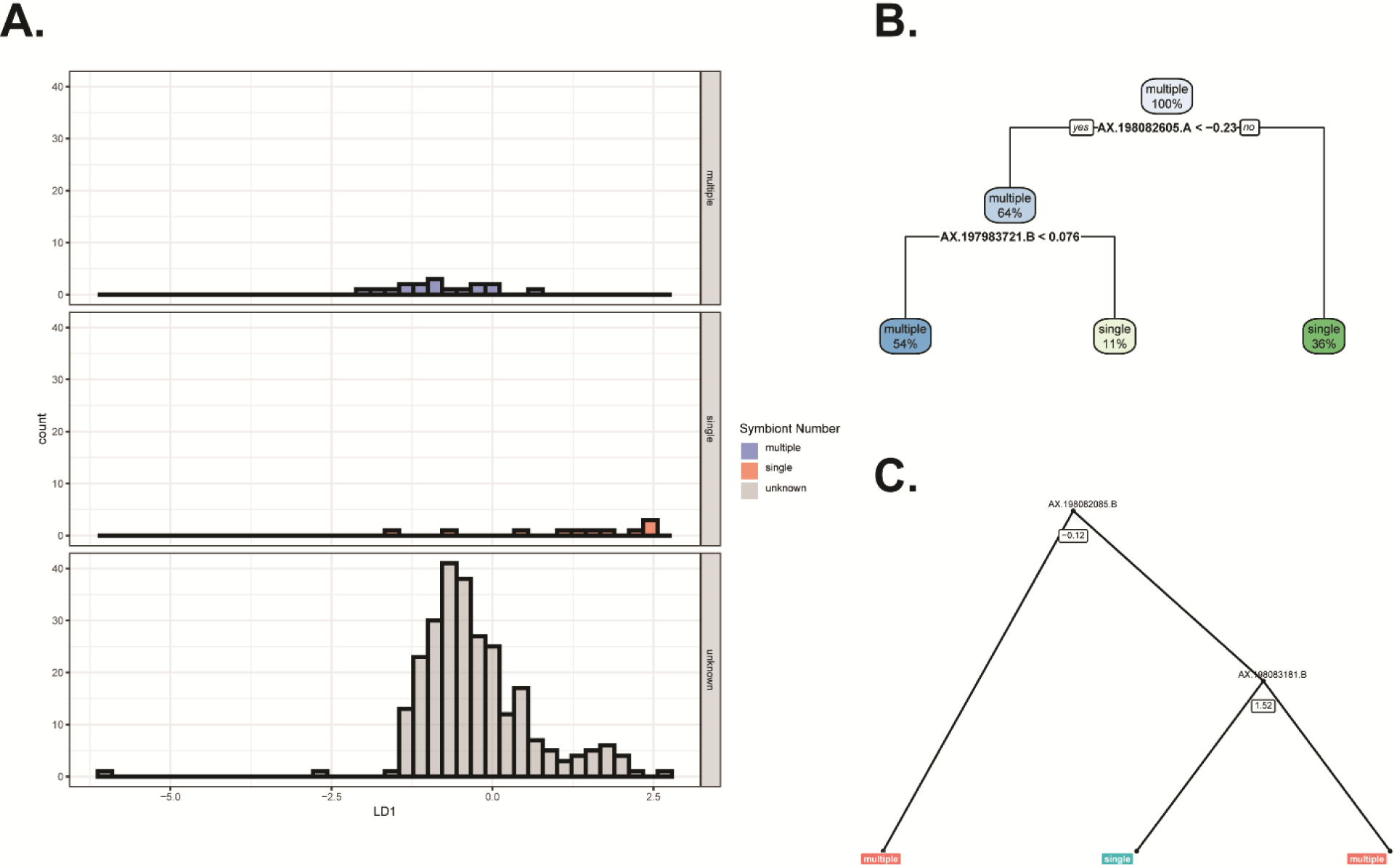
Single or multiple symbiont colonization. Linear discriminant analysis (A) decision tree (B) and random forest example tree (C).

**Figure S7.**
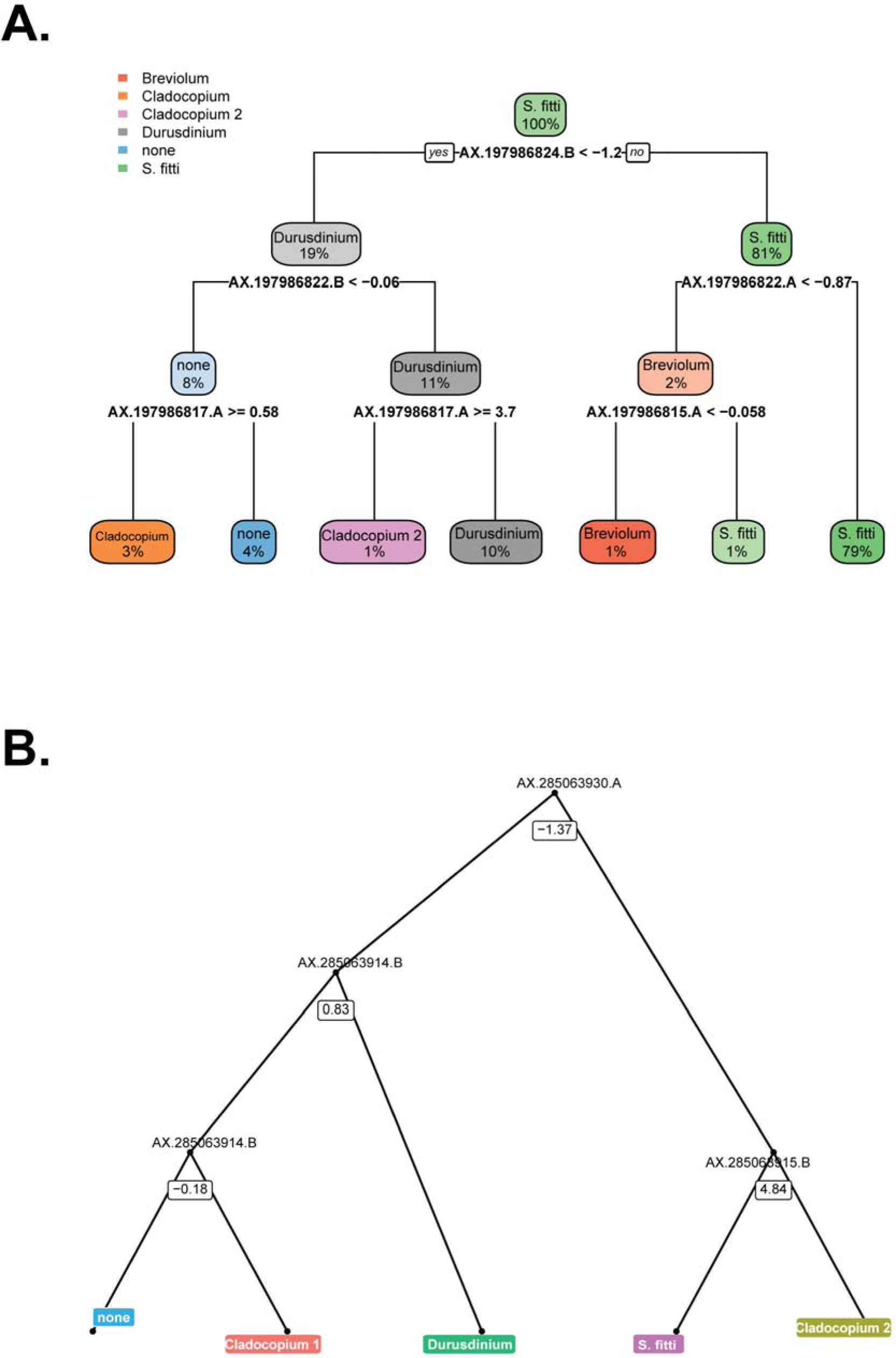
**Decision tree (A) and random forest tree (B) with the lowest error rate and maximum nodes for symbiont genera assignment.**

**Figure S8.**
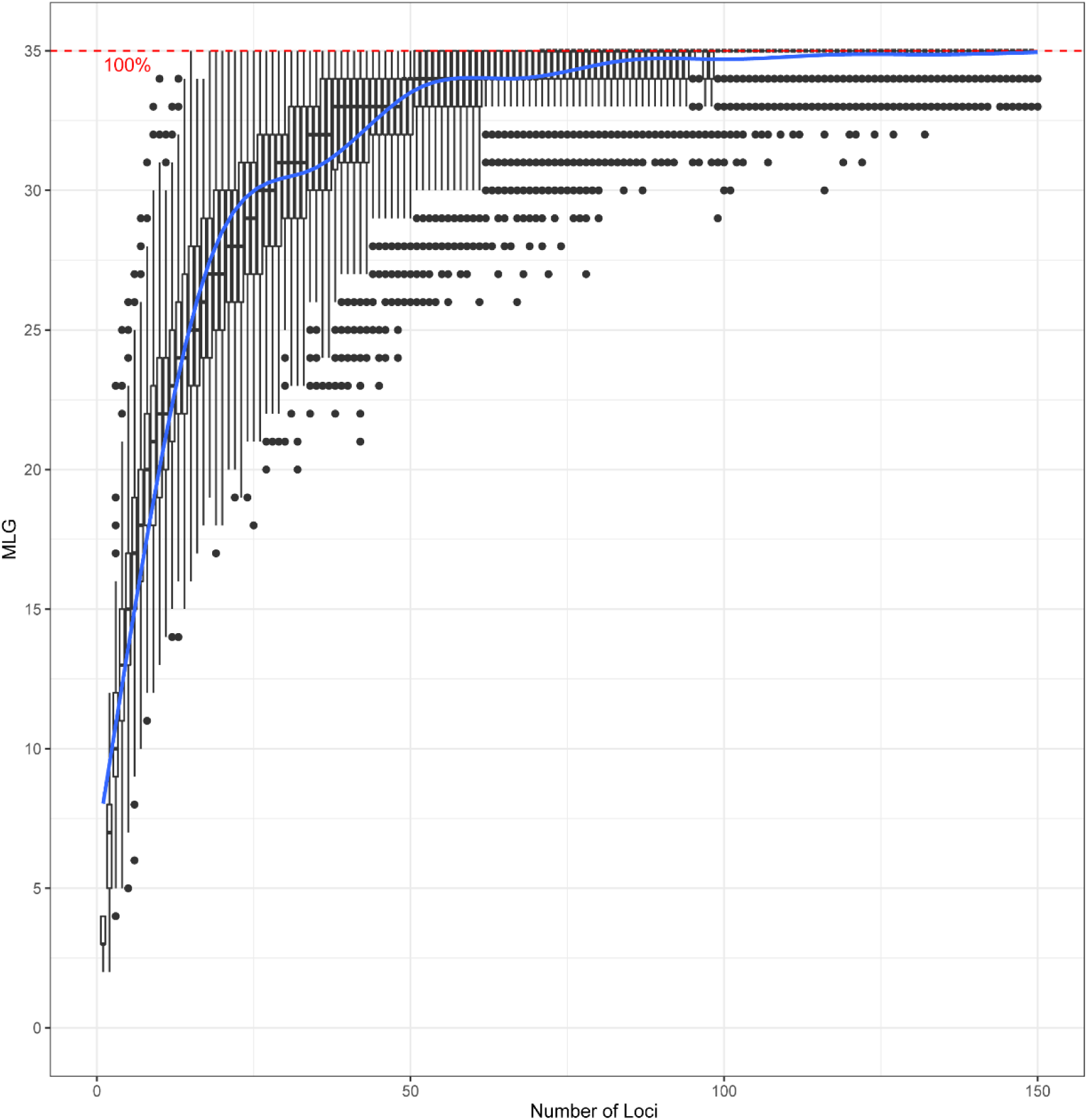
Genotype accumulation curve of the Pacific samples. The minimum number of loci required to recover 35 unique genet IDs. Boxplots are the results of the number of loci on the x-axis randomly sampled 100 times from all loci.

**Table S1.** Sample information for the five plates. (EXCEL)

**Table S2.**
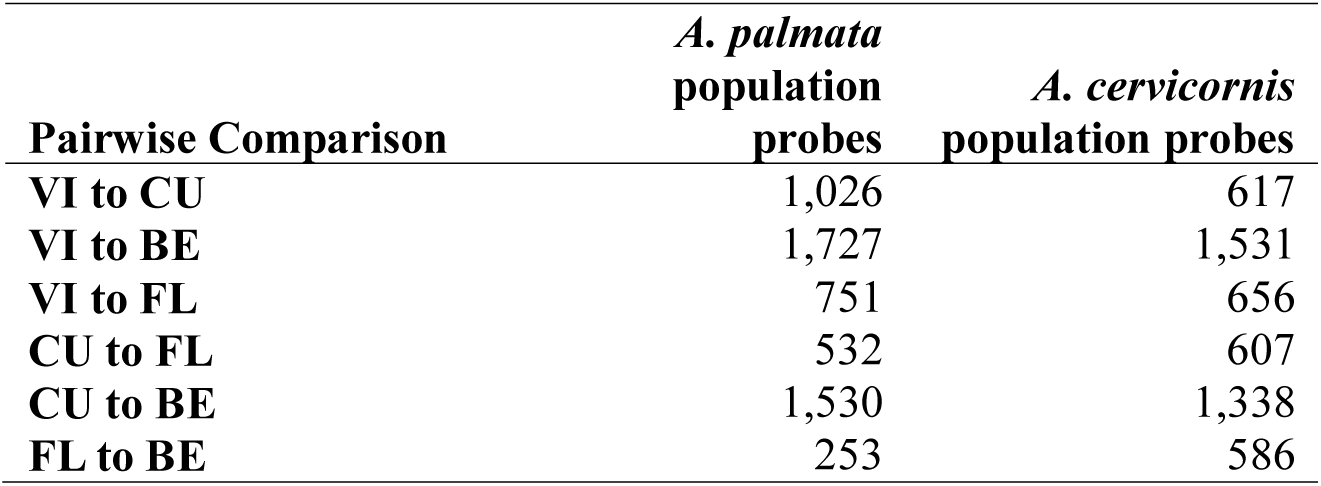
Number of population probes by species and location.

**Table S3.** **Sequence accession ID for genera probe design.**

| Gene | Accessions | References |
| --- | --- | --- |
| **cob** | JN557965.1, JN557953.1,<br>JN557943.1, JN557956.1,<br>JN557957.1 | (Pochon et al. 2012) |
| **COI** | JN557913.1, JN557901.1,<br>JN557891.1, JN557904.1,<br>JN557905.1 | (Pochon et al. 2012) |
| **cp23S** | JN558021.1, JN557991.1,<br>JN557969.1, JN558007.1,<br>JN558010.1 | (Pochon et al. 2012) |
| **elf2** | JN557889.1, JN557879.1,<br>JN557869.1, JN557882.1,<br>JN557883.1 | (Pochon et al. 2012) |
| **nr28S** | JN558091.1, JN558057.1,<br>JN558040.1, JN558075.1 | (Pochon et al. 2012) |
| **ITS2** | AF333507.1, AF333511.1,<br>AF499787.1, AF180124.1,<br>DQ480600.1, AF499793.1,<br>AF499797.1, AF334660.1,<br>Arif *et al.* ITS2 database | (LaJeunesse 2001; Reimer et al.<br>2006; LaJeunesse 2002; Arif et<br>al. 2014) |
| **psbA** | JN557866.1, JN557854.1,<br>JN557844.1, JN557857.1,<br>AB086863-AB086880.1 | (Pochon et al. 2012; Takishita et<br>al. 2003) |

**Table S4.**
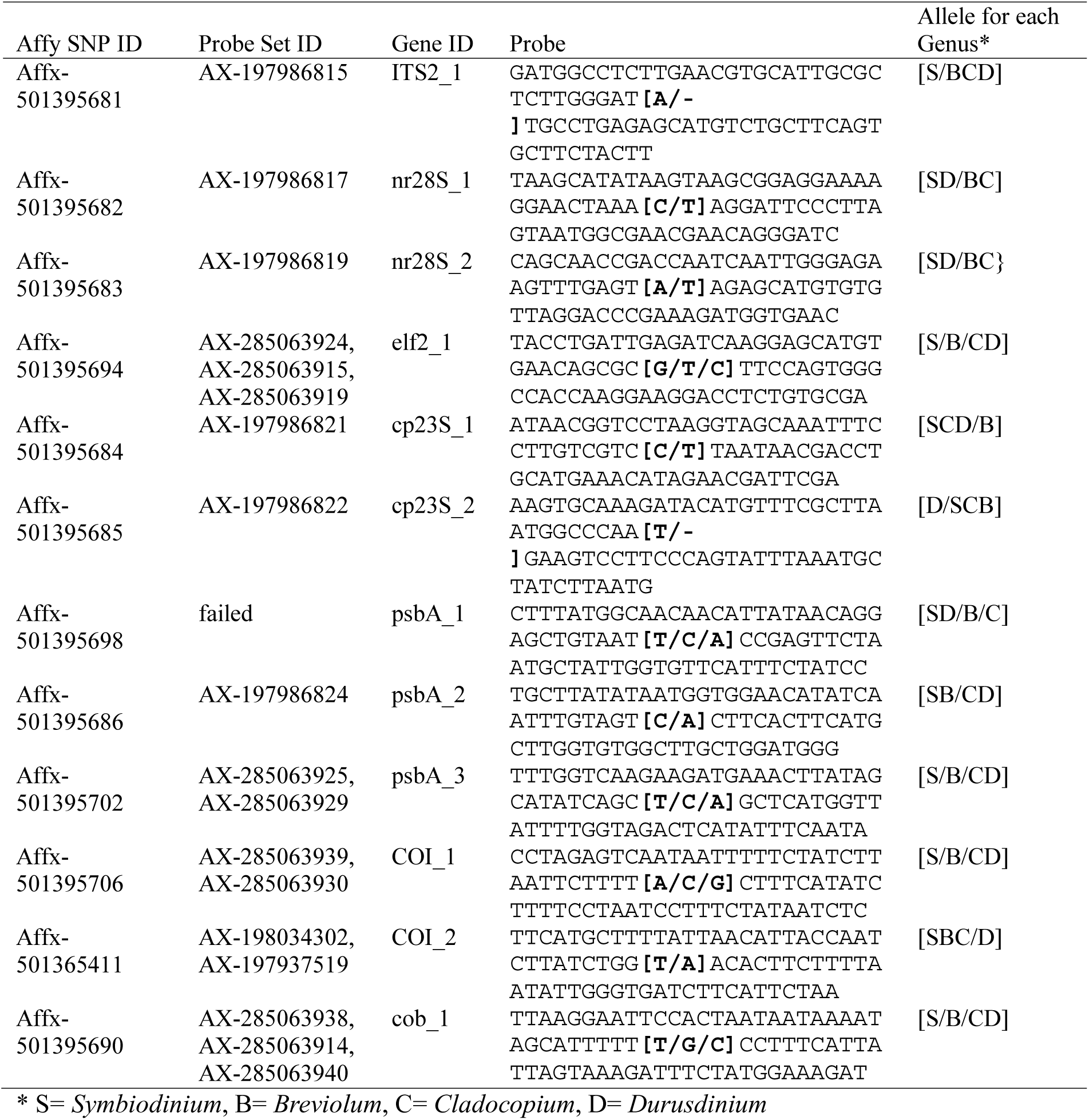
Symbiont genera probes.

**Table S5.** Caribbean Acropora recommended genotyping probe sets from five plates. (EXCEL)

**Table S6.** Symbiont recommended genotyping probe sets from five plates. (EXCEL)

**Table S7.** Pacific Acropora recommended genotyping probe set conserved across A. digitifera, A. millepora and A. muricata. (EXCEL)

**Table S8.** **Genetic distance between replicate DNA extractions and ramets of Sand Island samples.** Distances of DNA extractions originating from the same tissue sample are colored the same whereas distances between ramets are not colored.

| Sample ID | SI-1.1 | SI-1.2 | SI-1.3 | SI-10.1 | SI-10.2 | SI-10.3 | SI-12.1 | SI-12.2 | 11956 |
| --- | --- | --- | --- | --- | --- | --- | --- | --- | --- |
| SI-1.1 | 0 |  |  |  |  |  |  |  |  |
| SI-1.2 | 0.0041 | 0 |  |  |  |  |  |  |  |
| SI-1.3 | 0.0051 | 0.0049 | 0 |  |  |  |  |  |  |
| SI-10.1 | 0.0034 | 0.0047 | 0.0059 | 0 |  |  |  |  |  |
| SI-10.2 | 0.0071 | 0.0068 | 0.0061 | 0.0068 | 0 |  |  |  |  |
| SI-10.3 | 0.0020 | 0.0043 | 0.0050 | 0.0028 | 0.0063 | 0 |  |  |  |
| SI-12.1 | 0.0054 | 0.0054 | 0.0046 | 0.0049 | 0.0048 | 0.0042 | 0 |  |  |
| SI-12.2 | 0.0082 | 0.0067 | 0.0085 | 0.0076 | 0.0077 | 0.0078 | 0.0069 | 0 |  |
| 11956 | 0.0065 | 0.0093 | 0.0092 | 0.0071 | 0.0109 | 0.0060 | 0.0090 | 0.0123 | 0 |

**Table S9.** Genetic distance for each genotype with multiple ramets. (EXCEL)

**Table S10.** Reproducibility of genet identification between laboratories.

| User ID | Msat<br>geneti<br>ID | preserva<br>tive | Extraction<br>method | Lab | DNA<br>concentrati<br>on | SNP<br>genet ID | Missin<br>g Data | Heterozygosit<br>y | A.<br>cervicornis | A.<br>palmata | Genetic<br>Distanc<br>e |
| --- | --- | --- | --- | --- | --- | --- | --- | --- | --- | --- | --- |
| 13935 | C1522 | Ethanol | Qiagen<br>DNeasy | Baums | 34.71 | HG0136 | 1.44 | 1.06 | 97.66 | 0.22 | 0.020 |
| 13935_NF | C1522 | Ethanol | CHAOS | Fogarty | 0.064 | HG0136 | 1.96 | 6.01 | 91 | 0.21 |  |
| 3845 | C1548 | Ethanol | Qiagen<br>DNeasy | Baums | 22.76 | HG0003 | 0.36 | 0.57 | 98.81 | 0.3 | 0.018 |
| 3845_NF | C1548 | Ethanol | CHAOS | Fogarty | 1.894 | HG0003 | 1.49 | 5.71 | 91.96 | 0.22 |  |
| 13716 | C1639 | Ethanol | Qiagen<br>DNeasy | Baums | 16.97 | HG0006 | 1.23 | 2.04 | 96.51 | 0.29 | 0.027 |
| 126_NF | C1639 | CHAOS | CHAOS | Fogarty | 4.669 | HG0006 | 2.28 | 7.45 | 88.75 | 0.22 |  |
| 13829 | C1643 | Ethanol | Qiagen<br>DNeasy | Baums | 9.27 | HG0029 | 0.63 | 1.54 | 97.51 | 0.29 | 0.019 |
| 13829_NF | C1643 | Ethanol | CHAOS | Fogarty | 8.975 | HG0029 | 1.41 | 5.99 | 91.77 | 0.29 |  |
| 13839 | C1645 | Ethanol | Qiagen<br>DNeasy | Baums | 19.21 | HG0059 | 0.2 | 0.79 | 98.32 | 0.75 | 0.017 |
| 13839_NF | C1645 | Ethanol | CHAOS | Fogarty | 1.762 | HG0059 | 1.37 | 5.8 | 91.59 | 0.71 |  |
| 13923 | C1652 | Ethanol | Qiagen<br>DNeasy | Baums | 3.41 | HG0153 | 1.36 | 0.53 | 98.57 | 0.29 | 0.027 |
| 13923_NF | C1652 | Ethanol | CHAOS | Fogarty | 6.951 | HG0153 | 2.55 | 7.69 | 89.01 | 0.33 |  |
| 13756 | C1343 | Ethanol | Qiagen<br>DNeasy | Baums | 2.504 | HG0144 | 0.96 | 0.73 | 98.43 | 0.3 | 0.019 |
| 166_NF | C1343 | CHAOS | CHAOS | Fogarty | 4.951 | HG0144 | 1.78 | 6.28 | 90.89 | 0.29 |  |
| 1151 | P1020 | Ethanol | Qiagen<br>DNeasy | Baums | 22.16 | HG0171 | 1.08 | 1.3 | 0.3 | 97.59 | 0.012 |
| 1151_NF | P1020 | Ethanol | CHAOS | Fogarty | 8.154 | HG0171 | 1.42 | 3.32 | 0.3 | 94.64 |  |

## Notes

#### Summary of Updates

This version has been updated to fix grammatical errors and add additional permit information.

